# Biophysical neural adaptation mechanisms enable artificial neural networks to capture dynamic retinal computation

**DOI:** 10.1101/2023.06.20.545728

**Authors:** Saad Idrees, Michael B. Manookin, Fred Rieke, Greg D. Field, Joel Zylberberg

## Abstract

Adaptation is a universal aspect of neural systems that changes circuit computations to match prevailing inputs. These changes facilitate efficient encoding of sensory inputs while avoiding saturation. Conventional artificial neural networks (ANNs) have limited adaptive capabilities, hindering their ability to reliably predict neural output under dynamic input conditions. Can embedding neural adaptive mechanisms in ANNs improve their performance? To answer this question, we develop a new deep learning model of the retina that incorporates the biophysics of photoreceptor adaptation at the front-end of conventional convolutional neural networks (CNNs). These conventional CNNs build on ‘Deep Retina,’ a previously developed model of retinal ganglion cell (RGC) activity. CNNs that include this new photoreceptor layer outperform conventional CNN models at predicting primate and rat RGC responses to naturalistic stimuli that include dynamic local intensity changes and large changes in the ambient illumination. These improved predictions result directly from adaptation within the phototransduction cascade. This research underscores the potential of embedding models of neural adaptation in ANNs and using them to determine how neural circuits manage the complexities of encoding natural inputs that are dynamic and span a large range of light levels.

## Introduction

Artificial neural networks (ANNs) combined with deep learning algorithms are useful in modeling the function of the nervous system and are being used to model and investigate many brain areas (Doerig et al., 2023). Under relatively controlled conditions, ANNs perform well in computer vision tasks such as object recognition (Chollet, 2017; Simonyan and Zisserman, 2015; Krizhevsky et al., 2017), and they can successfully predict the responses of neurons in visual cortex (Kindel et al., 2019; Cadena et al., 2017; Yamins et al., 2014; Yamins and Dicarlo, 2016) and retina (McIntosh et al., 2016; Tanaka et al., 2019; Yan et al., 2022; Goldin et al., 2022; Maheswaranathan et al., 2018; Batty et al., 2017; Shah et al., 2022). However, it is less clear how ANNs perform in more naturalistic settings, where, for example, the statistics of sensory input can vary significantly from moment to moment. A specific concern is that the static nonlinear functions that ANNs typically employ will limit their ability to dynamically adapt to changing input conditions. This is important because adaptation is a nearly universal feature of individual neurons and neural circuits (Fain et al., 2001; Benda, 2021).

Sensory systems provide some of the clearest examples of the importance of adaptation: adaptation causes the fading of perceived intensity of odors (Zufall and Leinders-Zufall, 2000) and accommodation to sounds (Willmore and King, 2023). Within the visual system, adaptation constantly adjusts neural responses to match the prevailing input conditions. During natural vision, the amount of light falling on the retina can change locally and globally by several orders of magnitude on timescales ranging from fractions of a second (e.g. eye movements such as saccades), to minutes (e.g. movement between sunlight and shade), and to hours (e.g. the rising and setting sun). The limited dynamic range of individual neurons makes adaptation essential to match the range of neural responses to the current range of inputs. In vision, much of this adaptation occurs in the retina, and as early as the photoreceptors. Indeed, phototransduction can adapt rapidly and dynamically to control the sensitivity and kinetics with which light inputs are converted into electrical signals (Fain et al., 2001; Angueyra et al., 2022; Yu et al., 2022; Clark et al., 2013).

We examined whether incorporating photoreceptor adaptation into ANNs enhanced their accuracy at predicting neural responses under varied input conditions. We tested the ability of a convolutional neural network (CNN), similar to Deep Retina (McIntosh et al., 2016), to predict stimulus-evoked retinal ganglion cell (RGC) firing patterns under lighting conditions that differed from those under which they were trained. Observing a clear failure of the CNNs to generalize to new lighting conditions, we created a new type of CNN layer that incorporates a biophysical model of photoreceptor adaptation (Angueyra et al., 2022; Chen et al., 2024) that emulates the transformation of light into electrical signals. The photoreceptor layer can be used as an input to conventional CNNs, and can be trained end-to-end along with the other CNN layers. It thus equips deep learning CNN models of the retina with biorealistic adaptation mechanisms. We found these biophysical photoreceptor–CNN hybrid models better generalized across lighting conditions that were not included in training. Furthermore, because the photoreceptor adaptation was local, the photoreceptor–CNN model outperforms conventional CNN models in tasks involving rapid changes in local light intensity. These results suggest that chimeric models blending biophysical realism with trainable CNNs are better at modeling neural activity. Moreover they provide a promising direction for investigating how adaptation mechanisms shape neural circuit function.

## Results

### Photoreceptor adaptation improves CNN performance at predicting RGC responses to natural stimuli

We hypothesized that incorporating photoreceptor adaptation could improve the performance of CNN models at predicting RGC responses to naturalistic stimuli that involve local luminance variations. To test this hypothesis, we recorded the spiking activity of RGCs in isolated *ex vivo* primate (macaque monkey) retina using a high density multielectrode array (Methods), and then attempted to predict these visually-evoked responses using CNN models either with or without photoreceptor adaptation mechanisms. For training CNNs, we measured responses to a 36-minute checkerboard noise movie (30 *µ*m pixel edge) and to 9 naturalistic movies (totaling 9 minutes). The naturalistic movies were generated by displacing natural images from the Van Hateren dataset (Van Hateren and Van der Schaaf, 1998) across the retina, incorporating eye movement trajectories derived from the DOVES dataset (Van Der Linde et al., 2009) (Fig. 1a). The checkerboard noise provided responses to statistically stable stimuli, while the naturalistic movies presented large and rapid changes in light intensity characteristic of the retinal input during natural vision. Both stimuli were presented at a mean luminance of 50 R*receptor^*−*1^s^*−*1^, where rod photoreceptors are primarily driving RGC responses and rods are at a light level where their gain is adapting to the stimulus (Grimes et al., 2018; Griffis et al., 2023). We selected 57 RGCs (27 ON and 30 OFF parasol cells) for modeling purposes based on spike sorting quality and reliability across experimental conditions.

**Figure 1.**
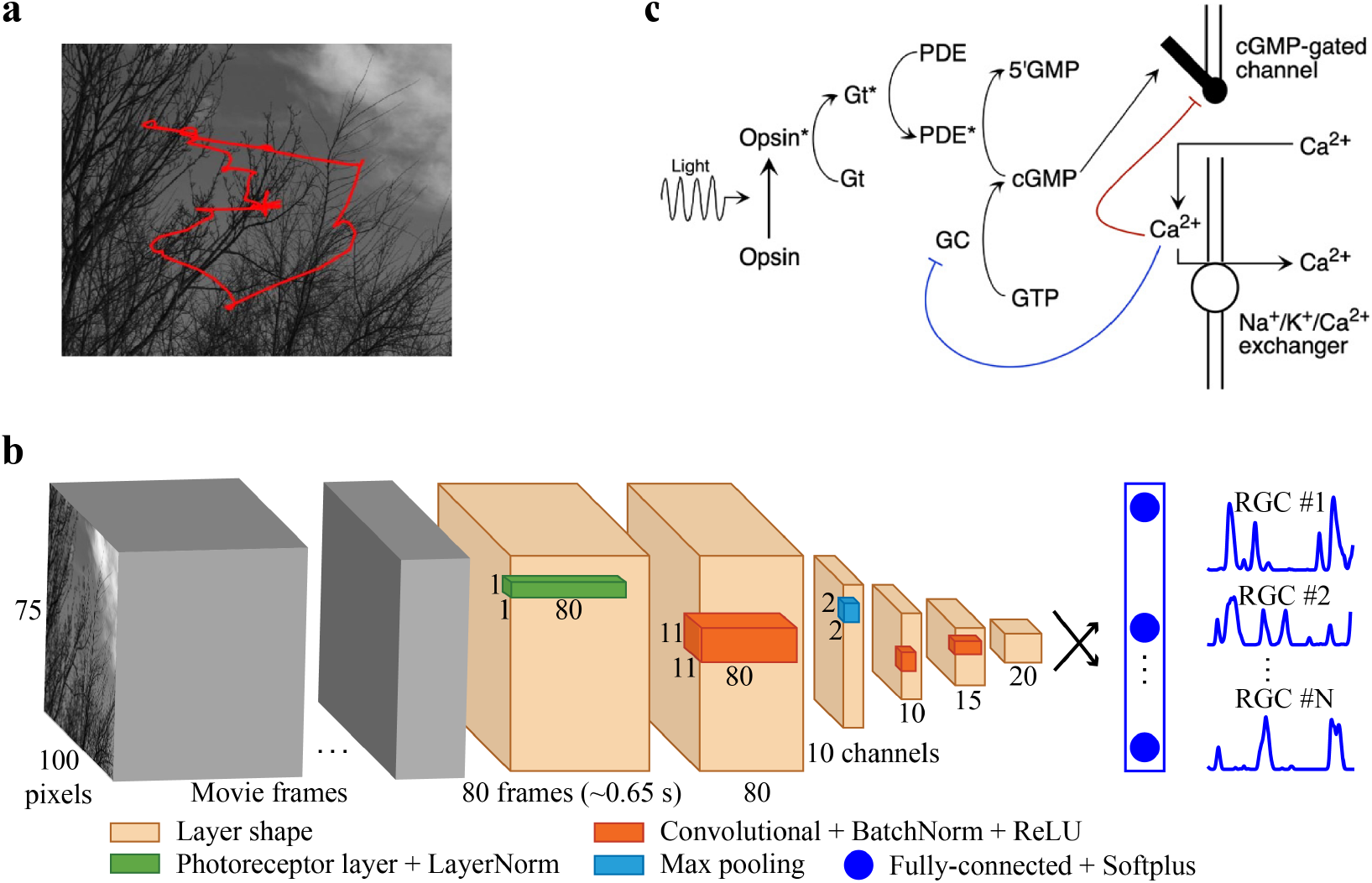
Training and architecture of photoreceptor–CNN / conventional CNN model. **a**. Naturalistic movie generated by displacing natural scene images across the retina, to mimic eye movement trajectories (red lines). **b**. Photoreceptor-CNN Model architecture incorporating a photoreceptor layer at the front-end (green) followed by Layer Normalization and 3 convolution layers (orange). The model output is a fully-connected layer that has N units based on the number of RGCs in the dataset followed by a softplus activation function that transforms the outputs into RGC spiking output (blue traces). By traversing through the input movie 80 frames at a time, an entire time series of RGC responses is obtained. When configured as a conventional CNN (without the photoreceptor layer), Layer Norm before the first convolution layer is the input layer. **c**. Schematic showing the phototransduction cascade and corresponding components of the biophysical model (reproduced from Angueyra et al. (2022)). Continuous synthesis of cGMP by guanylate cyclase (GC) opens cGMP-gated channels in the membrane. Activation of light-sensitive opsin (Opsin*) results in channel closure through the activation of G-protein transducin (Gt*), subsequently activating PDE* and decreasing cGMP concentration. Calcium ions (Ca^2+^) enter the photoreceptor outer segment via cGMP-gated channels and are extruded through Na^+^ / K^+^ / Ca^2+^ exchangers in the membrane. The biophysical model incorporates two distinct calcium-dependent feedback mechanisms influencing the rate of cGMP synthesis (blue line) and the activity of the cGMP-gated channels (red line). In the experiments presented here, the strength of Ca-dependent feedback to the the cGMP-gated channels was set to zero, based on observations from rod photoreceptor cells (Chen et al., 2024). See Supplementary Table. 1 for a list of parameters and their values.

We constructed a conventional CNN model similar to the existing state-of-the-art Deep Retina architecture (McIntosh et al., 2016) (i.e., the architecture in Fig. 1b with the photoreceptor layer removed, Methods). We re-optimized model hyperparameters – including the number of layers and the number of channels in each layer – for our dataset (see Methods). The resulting model had three CNN layers followed by a fully-connected layer with 10, 15 and 20 channels in each of the CNN layers, respectively, and filter pixel sizes (relative to the stimulus size) of 11x11, 7x7 and 7x7 respectively. Each convolution layer was followed by Batch Normalization (Batch Norm) followed by a rectifying nonlinearity. (Batch Norm z-scores the outputs of each unit in the CNN across the set of stimuli in each input batch.) The output of the fully-connected layer underwent a softplus operation to generate firing rates. This architecture resulted in 900,252 parameters, 873,642 of which were trainable. The model takes as input a movie segment of 80 consecutive frames, where each frame corresponds to 8 ms after upsampling the original 60 Hz frame rate to 120 Hz. The model output for the 80 consecutive frames provided an instantaneous spike rate for each RGC at the end of that movie segment. In other words, the response at each time point was based on the previous 80 frames (640 ms). By shifting the input forward by 1 frame at a time along the input movie, the model could predict the entire time series of RGC responses evoked by the stimulus movie. A Layer Normalization (Layer Norm) layer at the input standardized each pixel of the input stimulus segment relative to the values of that pixel in time. This normalization step removed the mean intensity from each input sample (movie segment) while preserving spatio-temporal contrast structure. This enabled the CNN to compensate for changes in input magnitude across light levels, ensured stability during training, and promoted faster convergence (Methods). Model performance was quantified using the fraction of explainable variance (FEV; see Methods) in the median-normalized response of each RGC (Methods) that was explained by the model. Henceforth, we present FEV as a percentage for ease of interpretation. A perfect model, by definition, would yield an FEV value of 100%. Comparisons between models were facilitated by presenting median FEV values along with 95% confidence intervals.

Due to the limited duration of the naturalistic movies in our dataset, we could not fully train the models on the naturalistic movie data alone. Instead, we adopted a two-stage training process. First, we trained the model to predict RGC responses using the entire white noise movie dataset, totaling 36 minutes. Second, we finetuned the resulting model using RGC responses to 8 of the 9 naturalistic movies (8 minutes of the data) and evaluated the model on the held-out movie (6 seconds). For an example RGC, this conventional CNN model captured 59% FEV of the recorded (Fig. 2a) response with a median FEV of 38% *±* 8% across the population (N = 57) of recorded cells (Fig. 3a).

**Figure 2.**
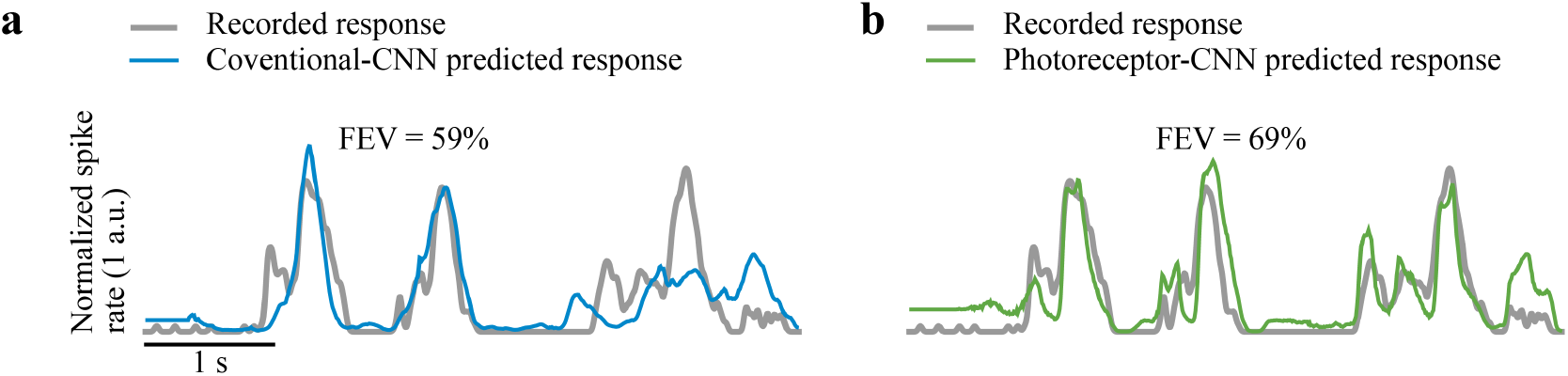
Incorporating photoreceptor adaptation improves CNN performance in predicting an example RGC’s response to naturalistic movies. **a-b**. Recorded response of an example primate parasol RGC to a held-out naturalistic movie shown as normalized spike rate (gray), and predicted responses by **(a)** conventional CNN model (blue) and **(b)** photoreceptor–CNN model (green). Fraction of Explainable Variance Explained (FEV) values quantify the percentage of variance in the RGC’s actual responses that could be explained by each model.

**Figure 3.**
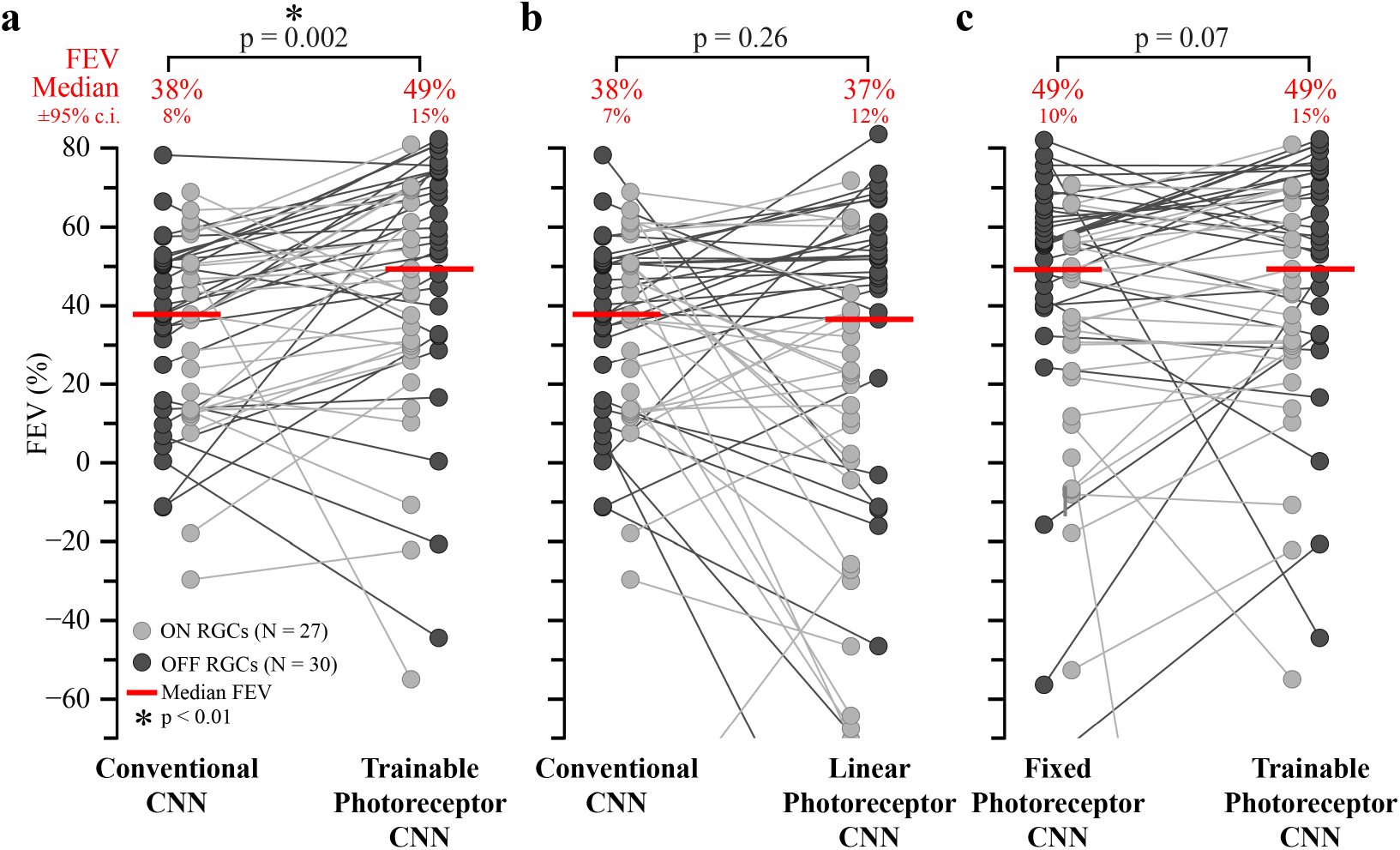
Incorporating photoreceptor adaptation improves CNN performance in predicting RGC responses to naturalistic movies. **a**. Comparison between conventional CNN and photoreceptor–CNN model with the photoreceptor layer parameters trained with the downstream CNN. Y-axis shows the performance of conventional CNN (left) and photoreceptor–CNN model (right) as FEV values for each RGC (circles). Light gray circles denote ON type RGCs (N = 27), and dark gray circles denote OFF type RGCs (N = 30). Connecting lines link the FEV values for each RGC across models. Median FEV values across all RGCs (N = 57) are indicated by red lines, and stated as FEV ^*±* 95% *c*.*i*.^ in red text at the top. P-values were calculated by performing two-sample Wilcoxon signed-rank test on the FEV distributions from the CNN and photoreceptor-CNN model. An asterisk indicates statistically significant difference (p < 0.01) between performance of the two models. **b**. Similar comparisons as in **a** but between conventional CNN (same as **a, left**) and photoreceptor-CNN, with the biophysical photoreceptor model being replaced by a linear empirical photoreceptor model. **c**. Similar comparison as in **a** between photoreceptor-CNN models with the biophysical photoreceptor layer parameters fixed to experimental fits to primate rods (left; Supplementary Table 1) and where the photoreceptor layer parameters were learned along with the downstream CNN (right; same as **a, right**).

To test the hypothesis that dynamic photoreceptor adaptation improves the CNN predictions, we developed a new type of CNN layer that builds upon a biophysical model of the phototransduction cascade by Angueyra et al. (2022). We then built a new CNN model that had this photoreceptor layer at its input and tested it with the same procedure used to test the conventional CNN (above). This photoreceptor model has previously been validated for its faithful representation of photoreceptor adaptation dynamics (Angueyra et al., 2022). The model is based directly on the signalling cascade that constitutes the phototransduction process (Fig. 1c). Rapid adaptation in this model emerges primarily from changes in the rate of cGMP turnover produced by light intensity-dependent changes in phosphodiesterase activity (Nikonov et al., 2000). The biochemical reactions of the phototransduction cascade were represented by a set of six differential equations that also incorporate dynamic feedback mechanisms to the cGMP-gated channels as detailed in the biophysical model by Angueyra et al. (2022) (reproduced in Supplementary Note 1). By setting the model’s parameter values to match experimentally-derived values from cone or rod photoreceptors (Chen et al. (2024); Supplementary Table. 1), the model can be configured to represent either photoreceptor type. Here, we configured the photoreceptor model to represent primate rods.

**Table 1.** Table of retinal reliability index for each retina used in this study.

| Retina | Retinal Reliability Index | Sorted RGCs |
| --- | --- | --- |
| Macaque retina 1: Natural stimuli experiments (Figs. 2, 3) | 0.90 | 57 |
| Macaque retina 2: Across light levels (Figs. 4, 5) | 0.96 | 37 |
| Rat retina 1: Across light levels (Figs. 7) | 0.99 | 61 |
| Rat retina 2: Across light levels (Figs. 7) | 0.99 | 58 |

The fully-trainable CNN layer encapsulating the biophysical photoreceptor model is characterized by twelve parameters (Methods) that could be trained together with the downstream network through backpropogation. This layer converts time-varying light intensity signals at each pixel in the input movie – measured in units of receptor activations per photoreceptor per second (R*receptor^*−*1^s^*−*1^) – into time-varying photocurrent values (measured in pA). In the hybrid biophysical photoreceptor–CNN model, the photoreceptor layer functions as the input stage of the CNN (Fig. 1b, photoreceptor layer). Following the photoreceptor layer, Layer Normalization is implemented to normalize the input distribution to the CNN, while preserving the spatiotemporal structure of the photocurrents. This design also ensures that the parameters of the biophysical model, having a different scale than the downstream CNN weights, can be trained together with the CNN through backpropogation. Henceforth, we refer to this hybrid model as the photoreceptor–CNN model. Unless explicitly stated, we allow the photoreceptor model parameters to be learned along with downstream CNN weights, and hence to deviate from the values to which they are initialized.

Similar to the conventional CNN, the photoreceptor– CNN model was trained to predict primate RGC responses first to a white noise movie and then parameters were fined tuned by training to naturalistic movies. When evaluated on the same held-out movie segment as the conventional CNN, the predicted responses of an example RGC generated by the photoreceptor–CNN model (Fig. 2b) were much more closely matched to the actual response than the predictions from the conventional CNN (Fig. 2b). Overall, the photoreceptor–CNN model performed with a median FEV of 49% *±* 15% (Fig. 3a). This performance gain is substantial given that the photoreceptor layer enhanced the predictive capability of conventional CNNs by approximately 29% (p = 0.002, two-tailed Wilcoxon signed-rank test, N = 57 RGCs). The enhanced capability of the photoreceptor– CNN model was robust when trained and evaluated over different combinations of naturalistic movies (Supplementary Fig. 1) and underscores the pivotal role played by the simulated photoreceptor layer.

The superior performance of the photoreceptor–CNN model can be attributed to its ability to capture and leverage dynamic photoreceptor adaptation. At the ambient light level of this experiment (50 R*receptor^*−*1^s^*−*1^) rod photoreceptors are adapting strongly (Grimes et al., 2018; Griffis et al., 2023) as sufficient numbers of photons are incident upon the rod outer segment to engage adaptation in phototransduction. Conventional CNNs, despite incorporating Layer Norm at their input to accommodate steady-state sensitivity changes associated with the mean intensity of the stimuli, struggle to capture this adaptation. Photoreceptor–CNNs explicitly model this dynamic adaptation, making them more effective in predicting RGC responses in this setting.

### Nonlinear adaptation in the photoreceptor layer drives performance gains

What causes the photoreceptor–CNN to outperform the conventional CNN at predicting RGC responses? Notably the superior performance is not attributable to an increase in model capacity from the addition of 12 trainable parameters of the photoreceptor layer. In fact, the photoreceptor–CNN model boasted a lower total number of trainable parameters (538,107 parameters) compared to the conventional CNN model (873,642 parameters). This difference in parameter count resulted from separately optimizing the hyperparameters of the photoreceptor–CNN and conventional CNN models via grid searches.

Given the inherent limitation of CNNs in dynamically adapting based on the prevailing inputs, we hypothesized that the observed performance gains in the photoreceptor–CNN model stem from the nonlinear dynamic adaptation and explicit feedback mechanisms embedded in the photoreceptor layer. To test this hypothesis, we substituted the biophysical photoreceptor model with an empirical linear photoreceptor model (Angueyra et al., 2022) (Methods). This linear model consists of a linear filter governed by 5 trainable parameters. These parameters were initialized to experimental estimates of single-photon responses obtained by measuring photoreceptor re-sponses (Angueyra et al., 2022). Unlike the biophysical model, the linear photoreceptor model lacks the ability to dynamically adjust its sensitivity. In this model, a single parameter provides sensitivity adjustment, applied to the entire linear photoreceptor model output to account for adaptation.

Following the same procedure as for the conventional CNN and the photoreceptor–CNN, the hyperparameters of the linear photoreceptor–CNN model were optimized via a grid search (Methods). The resulting model was first trained (including the 5 photoreceptor parameters that were initialized to experimental values) to predict RGC responses to the white noise movie and then finetuned using RGC responses to the naturalistic movies. When evaluated on the held-out segment of naturalistic movie data, this model performed very similarly to the conventional CNN model, yielding FEV of 37% *±* 12% (Fig. 3b; p = 0.26, two-tailed Wilcoxon signed-rank test, N = 57 RGCs). This is expected since the initial linear filtering stages of the conventional CNN should be able to capture the linear filtering performed by the simplified linear photoreceptor model.

Taken together, these experiments underscore the significance of nonlinear adaptation as crucial components influencing model predictions. Importantly, while the biophysical photoreceptor model parameters are trainable, similar performance gains could be achieved in another experiment where these parameters were set to non-trainable, thus reflecting true biological rods. In this experiment, the parameters of the photoreceptor layer were not learned or updated from their experimental fits to rods (Supplementary Table. 1) during the training of the photoreceptor–CNN model. Even with fixed parameter values, the resultant photoreceptor–CNN model exhibited performance comparable to its fully-trainable counterpart when evaluated on the naturalistic movie dataset (Fig. 3c; FEV 49% *±* 10%; p = 0.07, two-tailed Wilcoxon signed-rank test, N = 57 RGCs). These findings suggest that the observed performance gains can indeed be attributed to the nonlinear adaptation in the photoreceptors.

### Incorporating photoreceptor adaptation enables CNNs to generalize across light levels

The results so far indicate that the photoreceptor–CNN model shows superior performance in predicting responses to naturalistic movies that include dynamic local changes in intensity but lack changes in global light level. In the following sections, we shift our focus to asking how well the models generalize over global changes in mean light level, reminiscent of natural vision but occurring at slower time scales. This entails training the models at two distinct light levels and evaluating their performance on a third level not encountered during training.

To generate the experimental data for these computational studies, we recorded the spiking activity of RGCs in another isolated *ex vivo* primate (macaque monkey) retina using a high density multielectrode array (Methods). The retina was exposed to a binary checkerboard noise movie (Methods) with a checker size of 140 *µ*m, presented at three different light levels. The checkerboard pattern changed at 15 Hz and neutral density filters in the light path allowed for the mean intensity of the stimulus to be altered without changing the stimulus contrast. RGC responses were measured at mean intensities of 30 R*receptor^*−*1^s^*−*1^(high), 3 R*receptor^*−*1^s^*−*1^(medium) and 0.3 R*receptor^*−*1^s^*−*1^(low). Rod photoreceptors dominate signaling at all three light levels. At 30 R*receptor^*−*1^s^*−*1^the gain and kinetics of rod responses are adapting as sufficient numbers of photons are incident upon the rod outer segment to engage adaptation in phototransduction (Grimes et al., 2018; Griffis et al., 2023). However, at 0.3 R*receptor^*−*1^s^*−*1^rod responses are not adapting because individual rods are rarely receiving more than one photon at a time. RGC responses to the checkerboard noise were recorded for 60 minutes at each light level. The analysis focused on 37 RGCs based on spike sorting quality and reliably tracking cells across the light levels. These cells included 28 ON midget, 2 ON parasol, 2 OFF midget and 5 OFF parasol RGC types.

The conventional CNN architecture (same as the one used above) was re-optimized for the numbers of convolutional layers, filters in each layer, and the filter sizes with a grid search on this new dataset. The resulting model had three CNN layers followed by a dense layer with 8, 16 and 18 channels in each of the CNN layers, respectively, and filter sizes of 9x9, 7x7 and 5x5 respectively. The model takes as input an upsampled movie segment comprising of 120 consecutive frames, where each frame corresponds to 8 ms. We chose a larger segment length in these experiments to account for longer integration times at the lower light levels.

This CNN model was trained to predict RGC responses (N = 37 RGCs) (Methods) to a total of 40 minutes of a checkerboard noise movie presented at the high and medium light levels. To evaluate the model’s performance, we used a test data set that included 5–10 seconds of held-out segments of the checkerboard noise movie at all light levels (Fig. 4a), including the low testing light level not used during the training. Despite the same temporal sequence of checkerboard noise being presented at each light level, RGCs exhibited distinct responses, indicative of adaptation (Fig. 4b). The model accurately predicted responses to movies at the two training light levels (FEV of 93% and 83% for an example RGC, Fig. 4c, columns 1-2), with median FEVs of 84% *±* 11% for high and 78% *±* 3% for the medium light level (Fig. 5a). However, this model performed poorly at the low testing light level (Fig. 4c, column 3) with an FEV of only 24% *±* 15% across cells (Fig. 5a).

**Figure 4.**
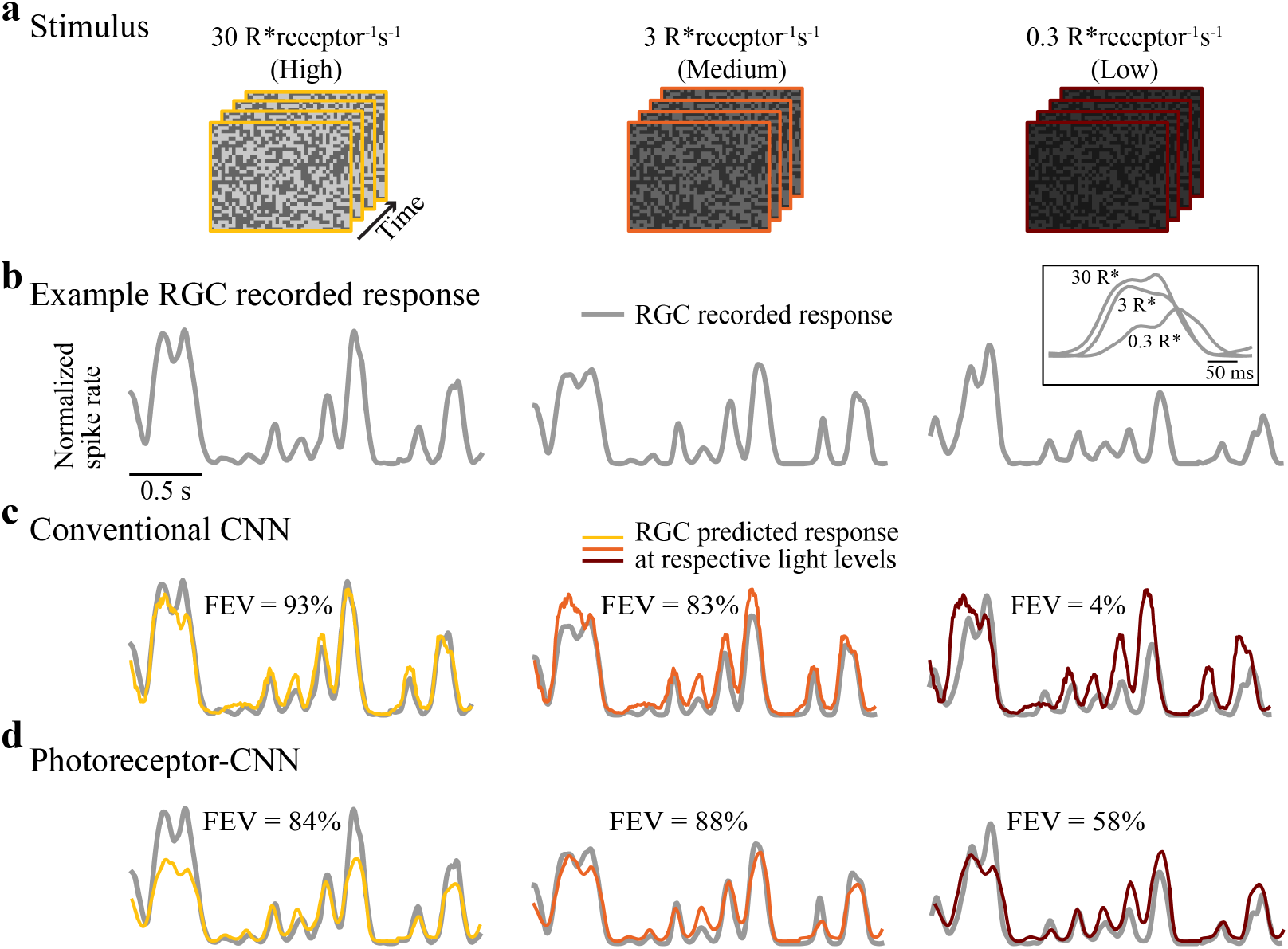
Incorporating photoreceptor adaptation enables CNN to predict responses of an example RGC at a light level different from those at which it was trained. **a**. White noise movie at three different light levels (columns). **b**. Recorded response (normalized spike rate) of an example RGC (gray lines) to white noise movie at the three different light levels in **a** (columns). Inset above the right column overlays a segment of the responses at the three light levels to directly compare response kinetics. **c**. Responses predicted by a conventional CNN model (colored) at each light level in **a** (columns). FEV values above each trace quantify the performance of the model for this RGC at the corresponding light levels. **d**. Same as in **c** but for the proposed photoreceptor–CNN model. Models were trained on data at high 30 R*receptor^*−*1^ s^*−*1^ (column 1) and medium 3 R*receptor^*−*1^ s^*−*1^ (column 2) and evaluated at low 0.3 R*receptor^*−*1^ s^*−*1^ (column 3) light level.

**Figure 5.**
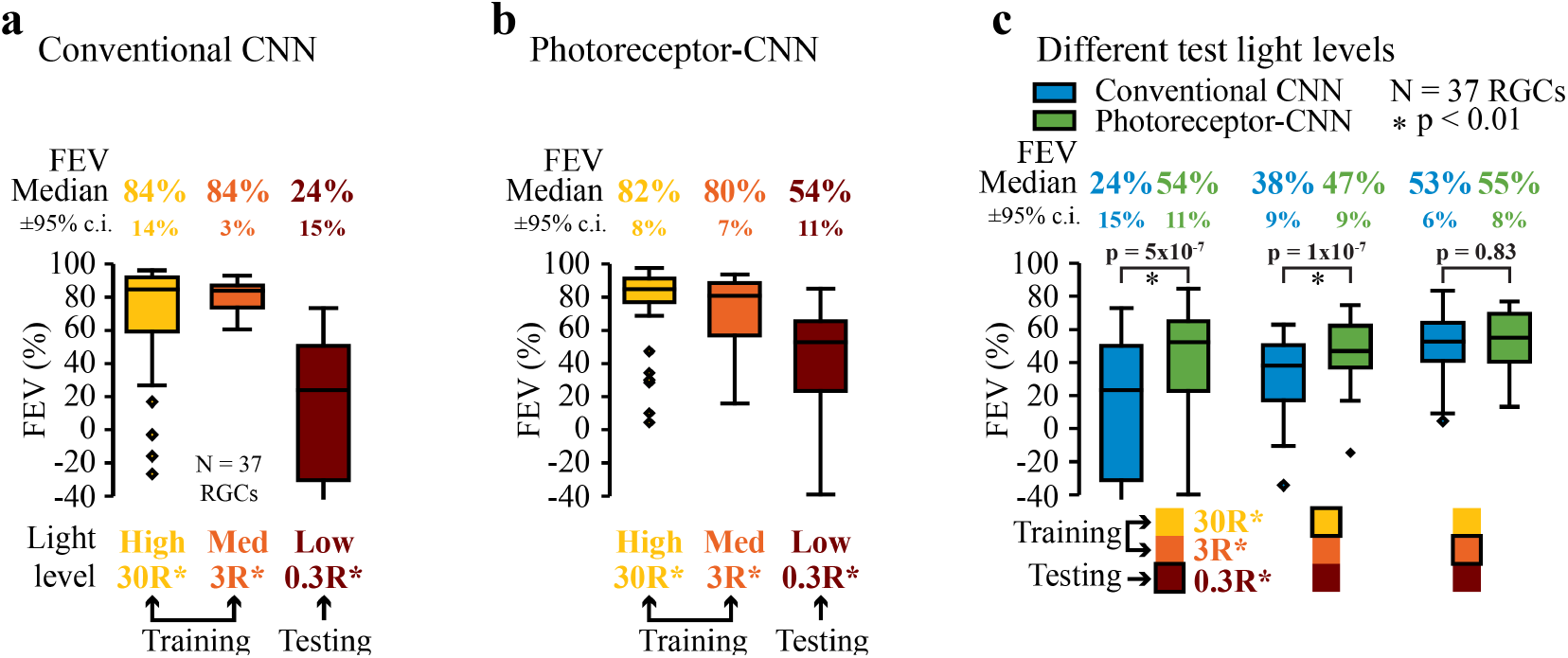
Incorporating photoreceptor adaptation enables CNNs to generalize across light levels. **a-b**. Performance of **(a)** a conventional CNN model, and **(b)** the photoreceptor–CNN model. Each model was evaluated at three light levels (labelled below each box plot): high (30 R*receptor^*−*1^ s^*−*1^) and medium(3 R*receptor^*−*1^ s^*−*1^), at which the models were trained, and low (0.3R*receptor^*−*1^ s^*−*1^) which the models did not see during the training. The box plot at each light level shows the distribution of FEVs across 37 primate RGCs. Numbers at the top of each box plot are the median FEVs ^*±* 95%*c*.*i*.^. **c**. Performance of the conventional CNN model (blue color; same model as in **a**), and the photoreceptor–CNN model (green color; same model as in **b**) at all combinations of training and test light levels. For each column, the legend below the box plot panel shows the two light levels the models were trained at and the third light level at which it was tested (black outline). The box plots show the distribution of FEVs at this testing light level. Testing light levels were low (column 1), high (column 2), and medium (column 3). p-values were calculated by performing two-sample Wilcoxon signed-rank test on the FEV distributions from the CNN and photoreceptor–CNN model at each testing light level.

Alterations in ambient light levels induce adaptation in the retina that alters both the sensitivity and kinetics of RGC responses (Tikidji-Hamburyan et al., 2015; Grimes et al., 2018; Ruda et al., 2020). CNN models can effectively capture linear sensitivity changes across global light levels, evident by consistent amplitudes of the predicted responses across light levels differing by a log unit (Fig. 4c). This is in part achieved by Layer Norm at the input to the model that discounts the mean intensity from input stimuli, resulting in a simple gain change commensurate with Weber adaptation. The failure of the CNN model at the low test light level (Fig. 5a, column 3) can therefore be primarily attributed to nonlinear sensitivity changes and shifts in response kinetics across the light levels, aspects not captured by Layer Norm alone. This is most prominent at the low test light level, where RGC responses slow considerably (Fig. 4b inset) and the inability of conventional CNNs to adaptively regulate both response sensitivity and kinetics limits their performance to generalize to this condition.

We next sought to determine whether incorporating the photoreceptor layer at the input stage could improve the ability of the CNNs to alter their sensitivity and kinetics in a light intensity-dependent manner and better predict the experimental data. To test this, we subjected the photoreceptor–CNN to the same test as the conventional CNN (above): we trained the model end-to-end (including the photoreceptor layer) to predict primate RGC responses to the binary checkerboard movie at the high and medium light levels. Similar to the conventional CNN model, the photoreceptor–CNN model reliably predicted responses to held-out stimuli at the two training light levels (Fig. 4d columns 1-2 and Fig. 5b). Importantly, the photoreceptor–CNN model could also explain 54% *±* 11% of the variance in responses at a light level lower than those at which it was trained (Fig. 4d column 3 and Fig. 5b). This performance was a two-fold improvement over the conventional CNN model without the photoreceptor layer (p = 5 × 10^−8^, Wilcoxon signed-rank test, N = 37 RGCs), which only explained 24% *±* 15% of the variance (Fig. 5a). We attribute this to the model’s ability to modulate output properties based on mean light level (see below).

Although we observed improved performance at the test light level, such gains were not evident at the training light levels (Fig. 5a,b). This is unlike our previously described experiments with the naturalistic movies (Fig. 3a) where the photoreceptor–CNN outperformed the conventional CNN even at the training light level. Noise stimuli have a more limited range of contrasts than natural scenes and lack temporal correlations. Hence, photoreceptor responses to noise stimuli at a single mean light level are nearly linear, with minimal changes in adaptation state. This leads to similar performance across conventional CNN and photoreceptor–CNN models (Fig. 5a,b) at the training light levels.

We also examined the performance of the photoreceptor–CNN across all different combinations of training and testing light levels: the model was trained on data from two light levels, and then evaluated on test data from a different light level (Fig. 5c; Supplementary Figure 2 shows the population data for ON and OFF RGCs separately). For all combinations of training and testing light levels, the photoreceptor–CNN model generalized better to new light levels than the conventional CNN model without the photoreceptor layer. The difference was smallest for the case where the testing light level (medium) was intermediate between the high and low training light levels (Fig. 5c, column 3; p = 0.83, two-tailed Wilcoxon signed-rank test, N = 37 RGCs). This is unsurprising given that conventional CNN models can interpolate between different sets of training conditions. Moreover, the similarity in responses at the high and medium light levels (inset in Fig. 4b) means that in the interpolation condition, the model predictions at the testing light level need not differ much from those at one of the training light levels. However, in the more challenging extrapolation tests – and especially in extrapolation to the low light level at which the response kinetics are appreciably different – the photoreceptor–CNN performs much better because the photoreceptor layer enables the models to adjust their output in a light-level-dependent manner.

### Photoreceptor-CNN model captures light-level-dependent changes in response kinetics

Normalization layers like Layer Norm allowed both the conventional CNN and photoreceptor–CNN models to capture steady-state sensitivity changes across light levels (Fig. 4c,d). But the inability of these normalization layers to adjust kinetics of the predicted responses suggested that the observed performance gains with the photoreceptor–CNN model resulted from its intrinsic ability to capture light-dependent changes in RGC response kinetics. For instance, RGC responses were faster (Fig. 6a) and temporal receptive fields had shorter latencies at the higher light level (Fig. 6b; estimated by reverse correlation). The response latency was consistently lower for the higher light level across the population of RGCs (Fig. 6e, left column). We then asked whether receptive fields of RGCs in the trained photoreceptor–CNN and conventional CNN models showed differences like those observed experimentally (Fig. 6b, Fig. 6e, left column). The temporal receptive fields of model RGCs (see Fig. 6c,d for an example RGC) were calculated by averaging the instantaneous temporal receptive fields (Supplementary Fig. 4a) across all movie segments. These instantaneous receptive fields were estimated by computing the gradients of the predicted RGC firing rates with respect to each input pixel value to yield instantaneous spatiotemporal receptive fields (Maheswaranathan et al., 2018; Goldin et al., 2022), which were then decomposed into spatial- and temporal components (Methods). We restricted this analysis to a subset of 22 RGCs that displayed FEV values greater than 50% at the training light levels (high and medium light levels) for both models (models of Figs. 4, 5a,b). Thus, these were the RGCs for which the predictions from both models were most reliable, facilitating estimates of receptive fields from model RGC gradients with respect to input stimuli.

**Figure 6.**
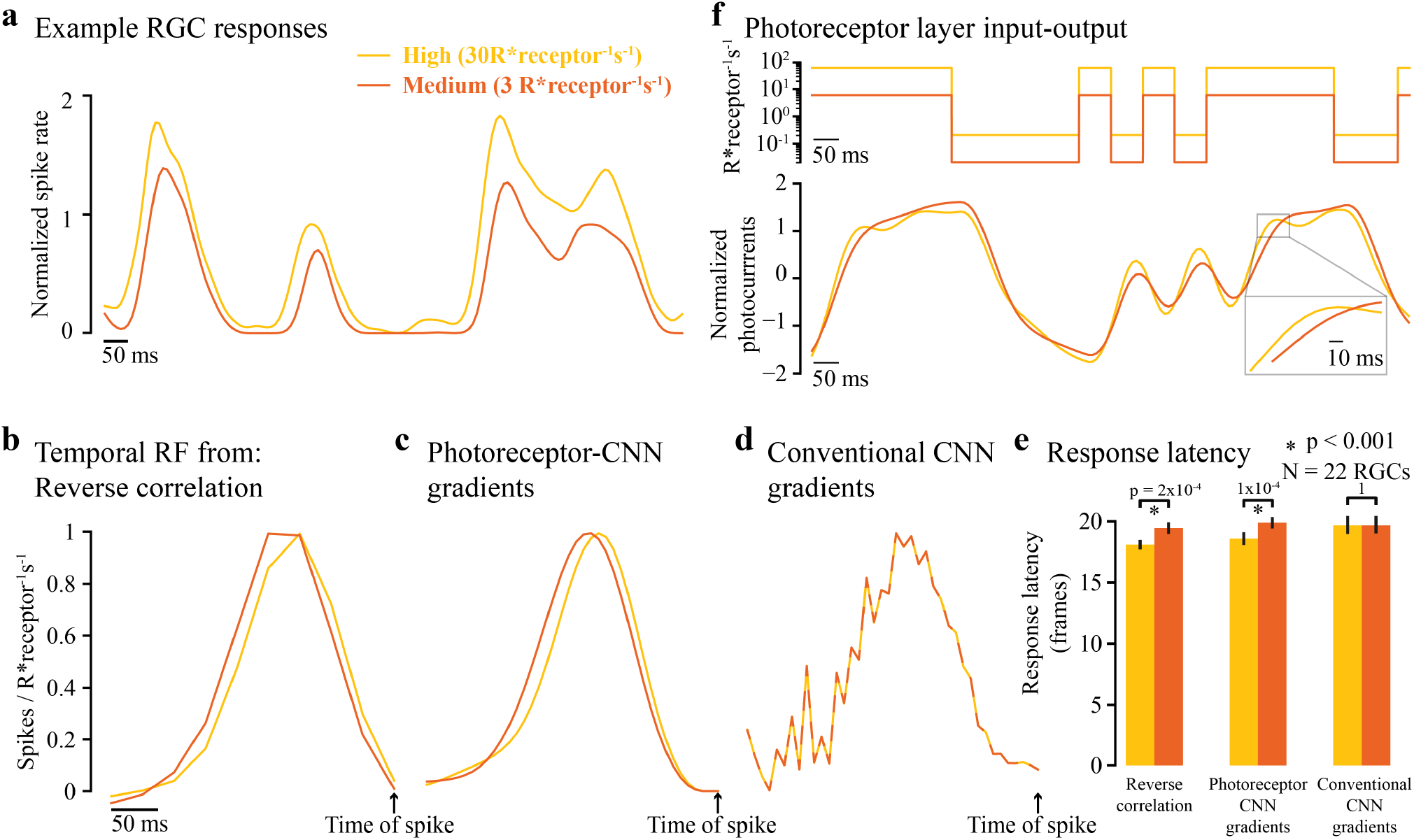
Photoreceptor layer enables CNNs to adjust their response kinetics in a light-level-dependent manner. **a**. Normalized recorded spiking activity of an example RGC in response to a white noise stimulus at two light levels used in model training: high (30 R*receptor^*−*1^ s^*−*1^; yellow), and medium (3 R*receptor^*−*1^ s^*−*1^; orange). **b**. Temporal receptive field of the same RGC calculated using reverse correlation of a white noise movie (55 minutes) at the two different light levels. **c**. Temporal receptive fields of the same RGC obtained by averaging across multiple instantaneous temporal receptive fields from photoreceptor–CNN output gradients with respect to multiple input movie segments. **d**. Same as in **c** but for the conventional CNN model. **e**. Mean response latency (N = 22 RGCs), calculated as the number of frames (1 frame = 8 ms) between the time of spike and peak of the temporal receptive field. Error bars indicate 95% confidence interval of the mean. Colors (legend in **a**) represent the light level at which temporal receptive fields were calculated from reverse correlation of experimental data (left column), photoreceptor– CNN model gradients (middle column) and conventional CNN model gradients (right column). An asterisk indicates a statistically significant difference in response latencies between two light levels (p < 0.001, N = 22 RGCs, two-tailed Wilcoxon rank-sum test). **f. Top**. Intensity changes over time for a single pixel in the binary checkerboard white noise movie at the two different mean light levels. **Bottom**. The output of the photoreceptor layer from the photoreceptor–CNN model after the Layer Normalization layer that immediately follows the photoreceptor layer. This output is fed into subsequent CNNs. Inset zooms the lag in photocurrents at medium light level (compare orange and yellow lines). Legend in **a** is valid for all panels (line style varies for clarity).

Consistent with our hypothesis, the temporal receptive fields of photoreceptor–CNN model RGCs exhibited intensity-dependent changes in response kinetics, with shorter latencies at the higher light level (Fig. 6c). This latency difference was statistically significant at the population level (Fig. 6e; p = 1 × 10^−4^, two-tailed Wilcoxon rank-sum test, N = 22 RGCs). This change in response latency could already be observed at the output of the photoreceptor layer (Fig. 6f), simply as a function of input light intensity. In contrast, the conventional CNN model showed no changes in latencies across the two light levels at which the models were trained (Fig. 6d; right column in Fig. 6e). This analysis underscores the effectiveness of the photoreceptor layer in capturing and adapting to dynamic changes in response kinetics associated with varying light conditions. In addition, photoreceptor–CNN models can also better capture sensitivity changes across light levels (see Supplementary Note 2).

### Incorporating photoreceptor adaptation enables generalization across photopic and scotopic light levels

Having observed that the photoreceptor–CNN model generalizes well across light levels that differ by 1-2 orders of magnitude (Fig. 5c), we wondered whether it could also generalize across more extreme variations in lighting. To answer this question, we trained conventional CNN and photoreceptor–CNN models to predict rat RGC responses to noise stimuli at a relatively bright photopic light level (10,000 R*receptor^*−*1^s^*−*1^) where cone photoreceptors predominantly contribute to vision, and evaluated the ability of the model to generalize to the much dimmer (scotopic) light level (1 R*receptor^*−*1^s^*−*1^) where rod (and not cone) photoreceptors are active (Fig. 7a). For this analysis, we used rat RGC recordings that we previously published (Ruda et al., 2020).

**Figure 7.**
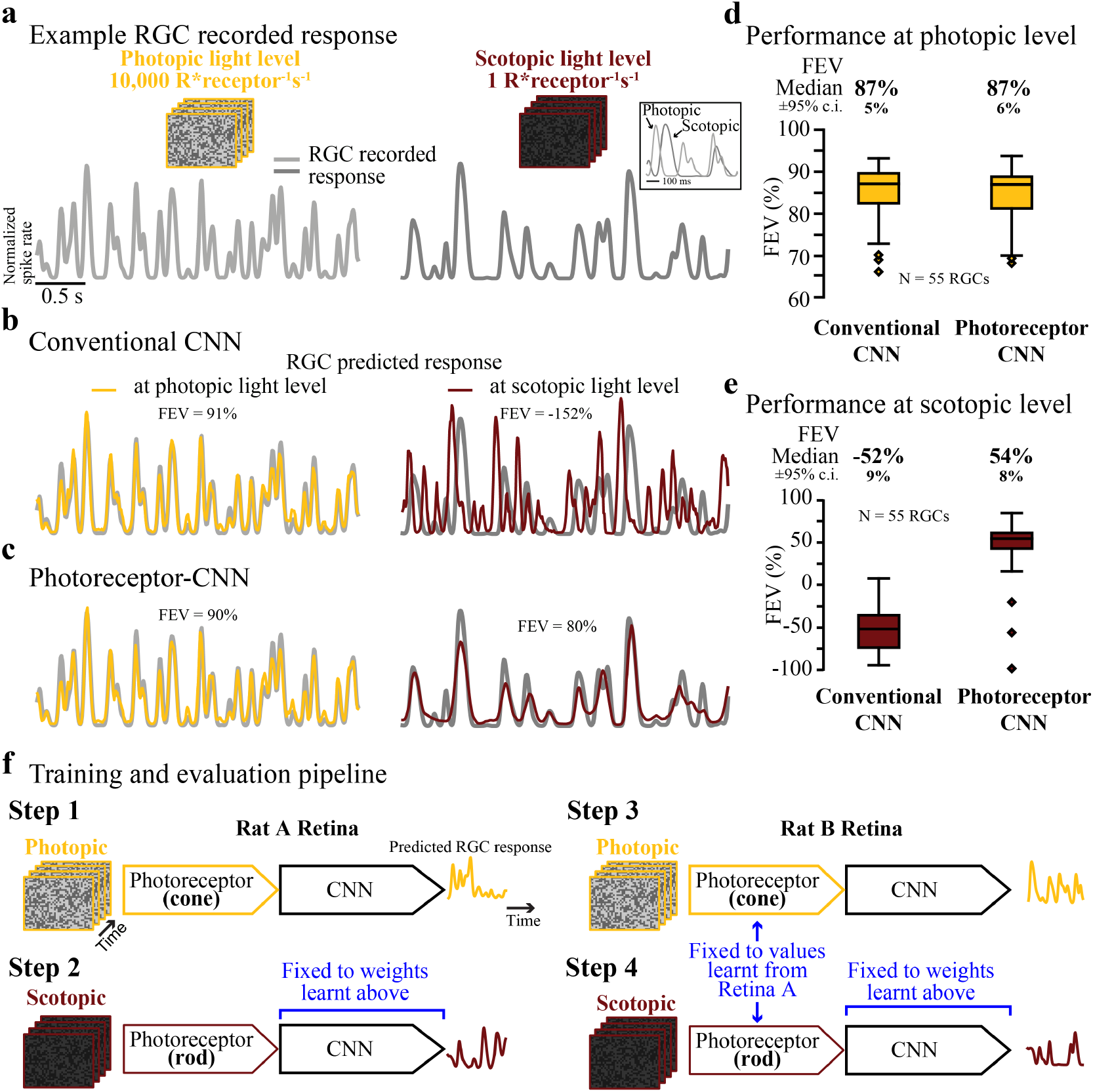
Incorporating photoreceptor adaptation enables CNN to generalize across extremely different light levels. **a**. Recorded responses shown as normalized spike rate (gray line) of Rat Retina B example RGC responses to held-out white noise at photopic light level (left) and scotopic light level (right). Inset shows an overlay of a segment of the responses at the two light levels. **b**. Responses predicted by a conventional CNN model (colored lines) at the two light levels in **a** when the model was trained at photopic light level only. FEV values above each trace quantifies the performance of the model for this RGC at the corresponding light levels. **c**. Same as in **b** but for the proposed photoreceptor–CNN model in which the CNN was only trained at photopic light level. The photoreceptor model was switched from cone to rod phototransduction model. **d-e**. Performance of conventional CNN model (left) and the photoreceptor–CNN model (right) when trained at photopic light level and evaluated at **(d)** photopic light level and **(e)** scotopic light level. The box plots shows the distribution of FEVs across RGCs (N = 55 RGCs, Retina B). Numbers at the top of each box plot are the median FEVs ^*±* 95%*c*.*i*.^. **f**. Schematic for training across extremely different light levels. **Step 1:** PR-CNN model was trained end-to-end to predict Rat Retina ‘A’ RGC responses at photopic light level. This led to an estimate for cone photoreceptor parameters and a model for the inner retina circuit (the CNN layers). **Step 2:** The model was re-trained at scotopic light level but the CNN layers were set to non-trainable and fixed to weights learnt in Step 1. In this case, the photoreceptor model learnt parameters reflecting rods. **Step 3:** The model was trained to predict Rat Retina ‘B’ responses at photopic light level. Photoreceptor layer was set to non-trainable with parameters fixed to cone parameters learnt in **Step 1. Step 4 (testing step):** Model was tested to predict Rat Retina ‘B’ responses at scotopic light level. Here, we used the rod photoreceptor parameters learnt in **Step 2** and rest of the model representing inner retina pathways of Retina B learnt in **Step 3**.

The conventional CNN model trained at the photopic light level could reliably predict RGC responses to heldout data at the photopic light level (example RGC responses in Fig. 7b, left; population data in Fig. 7d). However this model badly failed (FEV of *−*52% *±* 9%; N = 55 RGCs) to predict responses at the scotopic light level (Fig. 7b, right and Fig. 7e). The proposed photoreceptor–CNN model, however, did surprisingly well (Fig. 7c, right), achieving FEV of 54% *±* 8% on this task (Fig. 7e). For this experiment, we first trained the photoreceptor–CNN model at high light level and then replaced that model’s photoreceptor parameters (which correspond to cone cells at this light level) with those corresponding to rod cells (as explained in Fig. 7f and Supplementary Note 3). The remaining CNN parameters were unchanged by this procedure. This finding demonstrates that the changes in photoreceptor layer parameters alone can account for much of the difference in how the photoreceptor–CNN model predicts steady-state RGC responses at these two light levels.

## Discussion

We introduced a new CNN layer for vision models that builds upon a biophysical model of phototransduction (Angueyra et al., 2022). When used as a frontend to CNNs, this photoreceptor layer allows the CNN outputs to adapt to the prevailing inputs in a manner that more accurately mimics the retina. Consequently, the photoreceptor–CNN models surpass conventional CNN models at predicting RGC responses to naturalistic movies that simulate rapid local changes in light intensity due to eye movements, and at predicting responses across steady-state changes in mean light levels. The improved performance could not be replicated by replacing the biophysical photoreceptor model with a linearized photoreceptor model. Thus, the success of the biophysical photoreceptor–CNN model is attributable to nonlinear processes governing adaptation within the biophysical photoreceptor model.

ANNs, of which CNNs are a sub-class, are universal function approximators (Hornik et al., 1989) and therefore in principle they are capable of implementing any transformation with simple nonlinear units. This suggests that, *in principle*, a sufficiently large ANN can accurately model neural responses to stimuli with the same statistics as the training set, without the need for any bio-inspired adaptive mechanisms. Nonetheless, ANNs can benefit from having the right inductive biases that represent prior knowledge about the underlying data, as demonstrated by the benefits of CNNs in computer vision over non-convolutional forms of ANN (LeCun et al., 1989, 1998; Alzubaidi et al., 2021). In the same way, our results demonstrate that equipping CNN-based deep learning models with photoreceptor adaptive mechanisms improves their ability to capture retinal responses to stimuli with local luminance fluctuations, and enables them to better generalize to out-of-distribution tasks, such as extrapolating to new lighting conditions (Fig. 5c columns1-2, 7e). The new photoreceptor–CNN is much better at this challenging task as the photoreceptor layer enables the CNN to learn response properties such as the dependence of kinetics on light intensity (Dunn et al., 2006; Angueyra et al., 2022; Yu et al., 2022; Clark et al., 2013); Fig. 6c). This capability is demonstrated by the difference in kinetics of the responses at 3 R*receptor^*−*1^s^*−*1^and 30 R*receptor^*−*1^s^*−*1^(Fig. 6f, bottom); a fully linear model predicts identical kinetics at different mean light levels.

In addition to improving generalization between light levels, the photoreceptor–CNN model also substantially improves performance for predicting responses to high-resolution naturalistic stimuli at a single light level. Despite these improvements, the photoreceptor–CNN model is still far from a perfect predictor of retinal output. This is apparent in its inability to approach 100% FEV, or to achieve the same level of performance at held-out test light levels as it did at the light levels used for training (Fig. 5b). This limitation could be because the network lacks other adaptive mechanisms in the retina found downstream of the photoreceptors, such as spike frequency adaptation in RGCs (Chang et al., 2022; Brien et al., 2002). Introducing adaptive recurrent units (ARU; developed by (Geadah et al., 2022)) to the output layer of the photoreceptor-CNN, which implement spike frequency adaption through dynamic control of a nonlinearity, is a potential solution. ARUs at the output layer would also enable the network to have output units with diverse properties, similar to the diversity of RGCs (Wong et al., 2012; Goetz et al., 2022). Additionally, our current model does not explicitly capture the intricacies of adaptation in the intervening circuitry between photoreceptors and RGCs (i.e., in bipolar and amacrine cells). These include changes in gap junction coupling and switching between linear and nonlinear spatial summation (i.e., subunit rectification) (Grimes et al., 2014; Bloomfield et al., 1997; Bloomfield and Völgyi, 2004). To capture this adaptation, an adaptive-convolution layer based on a model for divisive gain control (Clark et al., 2013; Cui et al., 2016) could be used. This trainable layer would incorporate two pathways with distinct kinetics, with the output of one pathway controlling the sensitivity of the other, allowing for greater adaptability to changes in stimuli. Another limitation of the current approach is that Layer Norm was retained at the inputs of CNN in the photoreceptor–CNN model, normalizing the photoreceptor output. While this may inadvertently mitigate sensitivity changes across light levels, typically managed by the photoreceptor layer, it serves a crucial role in compensating for disparities in scale between the parameters of the biophysical model and the downstream CNN weights. The absence of Layer Norm negatively impacts the convergence of the photoreceptor–CNN model. Here, Layer Norm can, in principle, account for linear sensitivity changes linked to the mean intensity of a training sample (80-frame movie), but will have minimal impact on mitigating dynamic sensitivity changes triggered by fluctuations in pixel intensities within a training sample.

ANNs offer the potential to simulate networks of biological neurons, including those in the retina or visual cortex, making them highly relevant for visual neuroscience. These models are capable of automatically learning meaningful representations through multiple layers of abstraction. Establishing correspondence between ANN layers and neural layers (McIntosh et al., 2016; Maheswaranathan et al., 2018; Cowley et al., 2022; Yamins and Dicarlo, 2016; Mano et al., 2021) is increasingly providing insight into biological circuits. One challenge in using current ANNs to elucidate biological circuits is that ANNs are primarily designed to optimize performance on specific tasks rather than to mimic biological circuits. As a result, the structure and function of ANNs may not accurately reflect the complexity and organization of biological circuits which often include feedback loops and dynamic interactions between neurons. Moreover, ANNs typically consist of homogeneous units repeated throughout the network, which oversimplifies real neurons and neural circuitry and may fail to capture their full complexity. In contrast, the biophysical phototransduction model we use in the photoreceptor layer has parameters that map directly onto the biology, providing an opportunity for investigating the role of photoreceptor adaptation in the retina. For example, slower rod-mediated RGC responses at dim conditions compared to cone-mediated responses at brighter light levels (Ruda et al., 2020; Baylor and Fettiplace, 1977) may explain the temporal lag between predicted and actual response at dim light level (Fig. 7b, right): the conventional CNN model trained at bright light levels (10,000 R*receptor^*−*1^s^*−*1^) learned the faster kinetics of the cone pathway. While some of the differences in RGC responses may arise due to faster cone response kinetics (Cao et al., 2007; Baylor and Hodgkin, 1973; Ingram et al., 2016; Schneeweis and Schnapf, 1995), the relative contribution of photoreceptors and downstream retinal adaptation are not well understood. Fixing the photoreceptor layer parameters to empirically measured values, (like in Fig. 3c), can help in distinguishing photoreceptor from circuit mechanisms. Similarly, such biologically plausible models could provide insights into mechanisms underlying neural adaptation in other areas of the brain.

While our current findings indicate that the photoreceptor model we used offers superior performance (Fig. 3a) vs simpler photoreceptor models (like the linearized model shown in Fig. 3b), we acknowledge that other empirical models capturing similar dynamics may perform similarly well on the retinal prediction task. It is also possible that recurrent artificial units like the long short-term memory (LSTM) may partially capture photoreceptor adaptation effects by keeping track of arbitrary long-term dependencies in the input sequences. However, these units demand significantly more training data and introduce tens of thousands of parameters to the neural network. In stark contrast, the proposed biophysical photoreceptor layer only adds 12 parameters. Further, these parameters could be fixed based on direct photoreceptor recordings with minimal changes in CNN performance (Fig. 3c). In addition, the overarching goal is to integrate neural biophysics into ANNs to develop biologically interpretable computational models that surpass the limitations of conventional ANNs, offering a comprehensive framework for understanding complex biological phenomena.

In a similar vein, other neural predictor architectures, such as Generalized Linear Models (GLMs) could be used instead of CNNs, in conjunction with the photoreceptor model. Notably, we do not consider the CNN stages to be essential to our approach: rather, we consider the CNN to be the flexible scaffolding for incorporating the photoreceptor model into a trainable retina model. In future, this will allow incorporating fully-trainable biophysics models of downstream retinal components into this scaffolding. The result will be models with biologically-interpretable components that can predict retinal responses with high accuracy under varied conditions. We anticipate that these models could have substantial benefits for mechanistic investigations of visual function.

In general, models of retina that can leverage deep learning to model multiple ganglion cells simultaneously, together with biologically interpretable components, could be used to dissect the relative contributions by different cell types in the retina. They could also serve as an input stage to visual-cortical models to investigate higher visual processing under dynamic conditions. From a wider neuroscience perspective, this approach demonstrates the power of integrating neural dynamics in ANNs modeling brain functions where biophysical layers match the sensitivity to changing input conditions, while the downstream layers extract relevant features from dynamically adapting input stages. In summary, this approach establishes a framework to test which biological components are required to replicate brain function. Beyond neuroscience, these models could also pave the way for medical interventions, such as prosthetic devices that restore sight to the blind (Bosking et al., 2017).

## Methods

### Retina electrophysiology

Retina electrophysiology experiments were performed in two different labs. In the Manookin Lab, electrophysiological experiments were performed using ex vivo macaque retina (M. nemestrina) obtained through the tissue distribution program at the University of Washington National Primate Research Center and in accordance with the Institutional Animal Care and Use Committee at the University of Washington. Additional primate (Macaca mulatta) and rat retina electrophysiology experiments were performed in the Field Lab at Duke University, in accordance with Duke University’s Institutional Animal Care and Use Committee.

Electrophysiology experiments followed similar procedures in both the labs. For primate electrophyiology experiments, eyes were enucleated from terminally anesthetized macaque monkey and hemisected, and the vitreous humor was removed. Immediately after enucleation, the anterior portion of the eye and the vitreous were removed in room light. The eye cup was placed in a dark sealed container with Ames’ solution (Sigma, St. Louis, MO) at room temperature. Under infrared illumination, segments of peripheral retina 6-15 mm (25-70 deg, 200 *µ*m/deg) from the fovea and 3-5 mm in diameter were dissected and isolated from the retinal pigment epithelium. Preparation for the rat retinae was similar and described in detail in Ruda et al. (2020). Briefly, the retina of an euthanized animal was extracted and dissections were performed in darkness with the assistance of infrared converters. We dissected dorsal pieces of the retina that were 3 x 2 mm large. For recording, the retina was kept at 32-35^*°*^C and was perfused with Ames’ solution bubbled with 95% O_2_ and 5% CO_2_, pH 7.4.

The segment of retina was then placed flat, RGC layer down, on a planar multielectrode array (MEA) covering an area 2000 *µ*m x 1000 *µ*m. The MEA consisted of 512 electrodes with 60 *µ*m or 30 *µ*m spacing. Spikes on each electrode were identified by thresholding the voltage traces at 4 s.d. of a robust-estimate of the voltage s.d. For retina experiment involving naturalistic stimuli (Manookin Lab), spike sorting was performed using the Kilosort software package (version 2.5) Steinmetz et al. (2021). Spike waveform clusters were identified as neurons only if they exhibited a refractory period (1.5 ms) with *<*1% estimated contamination. For retina experiments across light levels (Field Lab), spike sorting was performed by an automated PCA algorithm and verified by hand with a custom software (Shlens et al., 2006; Field et al., 2007). Spike waveform clusters were identified as neurons only if they exhibited a refractory period (1.5 ms) with *<*10% estimated contamination. Most sorted units from the primate retina had 0% spike contamination based on refractory period violations. Other units had contamination in the range of 0.05%–0.09% with one unit at 0.8%. Only units that could be reliably tracked across all recording conditions were considered for further analysis.

For each retinal segment, a Retinal Reliability Index was computed to assess tissue quality. This involved analyzing the responses of individual sorted retinal ganglion cells (RGCs) to multiple trials using either white noise or naturalistic movies. We first estimated the trial-averaged noise by categorizing trials into two groups, averaging responses within each subgroup, and determining the mean squared error as

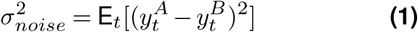

where, *y*^*A*^ and *y*^*B*^ are the observed spike rates of an RGC calculated as an average across set of trials *A* and set of trials *B* respectively. The sets *A* and *B* were obtained by randomly splitting the total number of repeats into two. We then computed the fraction of explainable variance which is the fraction of variance of each RGC attributable to the stimulus as

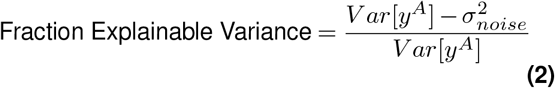

where,

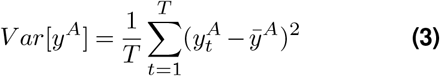

And 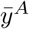 is the observed spike rate and *y*^*A*^ average across time. A higher fraction indicates recordings with low noise, where most of the variance in RGC responses is stimulus-driven. The retinal reliability index was determined by computing the median of the fraction of explainable variance across all sorted RGCs in each experiment. These values are presented in Table. 1 for each retina used in this study. Intuitively, higher the value (maximum 1), better the quality of recordings.

### Visual stimulation and data acquisition for primate retina experiment using naturalistic movies (Figs. 2–3)

Visual stimuli were created with custom Matlab code. Stimuli were presented with a gamma-corrected OLED display (Emagin, Santa Clara, CA) refreshing at 60.32 Hz. The display had a resolution of 800 x 600 pixels covering 3.0 x 2.3 mm on the retinal surface.

Spectral intensity profile (in *µ*Wcm^*−*2^nm^*−*1^) of the light stimuli was measured with a calibrated CCS100 spectrometer (Thorlabs). We transformed the stimulus intensity into equivalents of photoisomerizations per receptor per second (R*receptor^*−*1^s^*−*1^). The spectrum was converted to photons cm^*−*2^ s^*−*1^ nm^*−*1^, convolved with the normalized spectrum of macaque cones and rods, and multiplied with the effective collection area of these photoreceptors. The ambient light level (i.e. mean stimulus intensity) was set using neutral density filters in the light path. The attenuation of each neutral density filter was measured for the red, green, and blue LEDs using a calibrated UDT 268R radiometric sensor (Gamma Scientific).

We recorded RGC activity to 36-minutes of binary checkerboard white noise stimuli and 9-minutes of gray scale naturalistic movies at mean light level of 50R*receptor^*−*1^s^*−*1^. The checkerboard stimuli in this experiment had 100 x 75 pixels, where each pixel edge corresponded to 30 *µ*m on the retina surface. The refresh rate of the stimulus was set to 60.32 Hz (*∼* 16.6 ms per frame). The naturalistic movies were created by displacing natural scene images from Van Hateren dataset (Van Hateren and Van der Schaaf, 1998) across the retina, incorporating eye movement trajectories derived from the DOVES dataset (Van Der Linde et al., 2009) (Fig. 1a). We used 9 different natural scene images leading to 9 naturalistic movies. Each movie was 6-seconds long where the image remained stationary for the first 1-second to allow time to adapt to the spatial contrast before the motion began, which lasted for 5-seconds. The 9 movies were played in sequence and the entire sequence was repeated 10 times, totaling a duration of 9 minutes for naturalistic movies. These movies were presented to the retina at a resolution of 800 x 600 pixels where each pixel edge corresponded to approximately 3.8 *µ*m on the retina surface. We selected 57 RGCs (27 ON and 30 OFF parasol cells) for modeling purposes based on spike sorting quality and reliability across experimental conditions.

At the model training stage, each repeat of the movie was treated as an individual movie, i.e., 9 movies with 10 trials were treated as 90 movies. 80 of which (8 unique movies and 10 trials) were used for training the model and the held-out movie was used to validate the model against trial averaged responses. Additionally, naturalistic movies were spatially down-sampled by a factor of 8 to 100 x 75 pixels to match the resolution of checkerboard white noise stimuli. This was necessary as we first trained the models on checkerboard movie and then fine-tuned the same model with naturalistic movies.

### Visual stimulation and data acquisition for primate and rat retina experiment at different light levels (Figs. 4–7)

Visual stimuli were created with custom Matlab code. Stimuli were presented with a gamma-corrected OLED display (SVGA + XL Rev3, Emagin, Santa Clara, CA) refreshing at 60.35 Hz. The image from the display was focused onto the photoreceptors using an inverted microscope (Ti-E, Nikon Instruments) with a x4 objective (CFI Super Fluor x4, Nikon Instruments). The optimal focus was confirmed by presenting a high spatial resolution checkerboard noise stimulus (20 x 20 *µ*m, refreshing at 15 Hz) and adjusting the focus to maximize the spike rate of RGCs over the MEA. The display had a resolution of 800 x 600 pixels covering 4 x 3 mm on the retinal surface.

Spectral intensity profile (in *µ*Wcm^*−*2^nm^*−*1^) of the light stimuli was measured with a calibrated Thorlabs spectrophotometer (CCS100). We transformed the stimulus intensity into equivalents R*receptor^*−*1^s^*−*1^by converting the power and the emission spectra of the display to an equivalent photon flux by Planck’s equation. This converted the emission spectrum to photons cm^*−*2^ s^*−*1^ nm^*−*1^, which was then convolved with the normalized spectral sensitive of rods Baylor et al. (1984), and multiplied with the effective collection area of rods (0.5 *µ*m^2^). The ambient light level (i.e. mean stimulus intensity) was set using neutral density filters in the light path. In each recording, stimuli were first presented at the darker light level. For every subsequent higher light level, the retina tissue was first adapted to that light level before continuing the recordings.

Stimuli consisted of non-repeated, binary checkerboard white noise interleaved with repeated (N = 126 or 225 trials), binary white noise segments (5 or 10 s) to estimate noise. The total duration of stimulation was 60 minutes. We recorded primate RGC activity to the same 60 minute white noise sequence at three different mean light levels, each differing by 1 log unit: 0.3 R*receptor^*−*1^s^*−*1^, 3 R*receptor^*−*1^s^*−*1^and 30 R*receptor^*−*1^s^*−*1^. These light levels fall under the scotopic regime, where mostly rod photoreceptors contribute to vision. The movies across the three light levels only differed in their mean pixel values which were 0.3 R*receptor^*−*1^s^*−*1^(low light level), 3 R*receptor^*−*1^s^*−*1^(medium) and 30 R*receptor^*−*1^s^*−*1^(high). The checkerboard stimuli in this experiment had 39 pixels x 30 pixels, where each pixel edge corresponded to approximately 140 *µ*m on the retina surface. The refresh rate of the stimulus was set to 15 Hz which means that each checkerboard pattern was exposed on to the retina for *∼* 67 ms. In this work, we used a subset of 37 recorded RGCs that could be reliably tracked across light levels and were classified as high quality units after spike sorting. This subset contained 2 ON parasol, 28 ON midget, 5 OFF parasol and 2 OFF midget RGC types).

The rat experiments of Ruda et al. (2020) were performed at two light levels differing by 4 log units: 1 R*receptor^*−*1^s^*−*1^(scotopic light level where mostly rod photoreceptors contribute to vision) and 10,000 R*receptor^*−*1^s^*−*1^(photopic light level where cone photoreceptors predominantly contribute to vision). The white noise checkerboard movie in these experiments had 10 pixels x 11 pixels, with each pixel edge corresponding to approximately 252 *µ*m on the retina. The refresh rate of the stimulus was 60 Hz and 30 Hz at the photopic and scotopic light levels, respectively. In this work, we used data from two rat experiments: a subset of 61 RGCs from Retina A and a subset of 55 RGCs from Retina B that could be reliably tracked across light levels and were classified as high quality units after spike sorting. This subset contained OFF brisk sustained and OFF brisk transient RGC subtypes.

### Data preparation for models

Both white noise and naturalistic movies were upsampled to 120Hz by repeating each frame so that each frame had a duration of 8 ms. This up-sampling was necessary for the photoreceptor layer in which differential equations are solved using the Euler method.

Spikes were grouped in 8 ms time bins spanning the duration of the movie. Firing rates were then estimated by convolving the binned spike counts with a Gaussian of *σ* = 32 ms (4 frames/bins) standard deviation and amplitude of 0.25*σ*^*−*1^*e*^1*/*2^. The resulting firing rates for each RGC were normalized by the median firing rate of that RGC over the course of the experiment. This was done to ensure that responses of all output units of the model (i.e. the modeled RGCs) were at the same scale.

### Conventional CNN architecture

The general architecture of the conventional convolutional neural network (CNN) we used was similar to Deep Retina (McIntosh et al., 2016). The model (Fig. 1b) had 3 convolution layers (orange color), followed by a fully-connected output layer (black arrows). The model takes as input a movie (80 frames per training example where each frame corresponds to 8 ms) and outputs an instantaneous spike rate for each RGC at the end of that movie segment. The first convolution layer is a 3D convolutional layer operating in both the spatial and temporal dimensions. The output of the 3D convolutional layer is a 2D image which is normalized using Batch Normalization (BatchNorm) and then passed through a rectifying (ReLU) nonlinearity. All the temporal information from the movie is extracted by this layer as the temporal dimension of the convolutional filter is the same as the temporal dimension of the movie. To down sample the spatial dimensions, we applied a 2D max pool operation (blue color) that took the maximum value over 2x2 patches of the previous layer’s output. The subsequent 2D CNN layers are followed by a final, fully connected layer with softplus activation function that outputs the predicted spike rate for each RGC in the dataset.

To obtain the time series of RGC responses to longer movie stimuli, we feed into the model many 80-frame video samples taken from that longer movie, that correspond to 1-frame shifts. I.e., the model receives as inputs frames 1–80, 2–81, 3–82, etc., and outputs RGC responses at the times of movie frames 80, 81, 82, etc.

A Layer Normalization (Layer Norm) at the input of the first convolutional layer was applied to z-score each frame of a movie segment. Layer Norm computes normalization statistics for each pixel over its temporal history within a single training example i.e. a single movie segment comprising 80 frames. This step removes the mean luminance from each training example, mitigating sensitivity changes associated with global luminance changes while preserving spatio-temporal structure within each movie segment.

Each convolution operation is followed by Batch Normalization which contributes to stable training and faster convergence of the model (Ioffe and Szegedy, 2015). During model training, the distribution of inputs to a layer undergoes changes as the network’s parameters are updated, leading to what is known as the internal covariate shift – a phenomenon that hampers model convergence and introduces instabilities. These Batch Norm layers address the internal covariate shift during training by z-scoring the input, using normalization statistics computed based on batch statistics from batches comprising over 100 movie segments. This process enforces a 0 mean and unit variance for the data, introducing two trainable parameters – shift and scale – that systematically adjust weights and biases in the CNN layer. Additionally, the running average and variance of the training data serve as non-trainable parameters saved for later use during the test phase. This normalization process mitigates extreme parameter values, preventing issues such as exploding or vanishing gradients. The scale and shift parameters enable the model to adapt to variations in feature magnitudes across layers and activations, facilitating improved and faster convergence. During the test phase, Batch Norm uses the non-trainable moving average and variance saved during the training to normalize its inputs i.e., the outputs of the convolutional layer. Batch Norm then scales and shifts the normalized input using the scale and shift parameters learned during the training phase. Notably, in the current setup, Batch Norm parameters are not influenced by the mean light level as Layer Norm at the model’s input removes mean luminance from each training sample.

For modeling RGC responses across light levels (experiments of Figs. 4–7), the input to the model was a movie segment of 120 frames instead of 80 frames. The longer movie segment allowed for longer integration times at the lowest light level of 0.3 R*receptor^*−*1^s^*−*1^.

### Biophysical photoreceptor–CNN architecture

The proposed photoreceptor convolution layer builds upon a biophysical model of the phototransduction cascade by Angueyra et al. (2022) (Fig. 1c). The model incorporates the various feedforward and feedback molecular processes that convert photons into electrical signals, and therefore faithfully captures the photoreceptors’ adaptation mechanisms. The biophysical model is reproduced in Supplementary Note 1 and described in brief below.

The biophysical model was represented by a set of six differential equations that mimics the enzymatic reactions of the phototransduction cascade. Rapid adaptation in this model emerges from changes in the rate of cGMP turnover produced by light intensity-dependent changes in phosphodiesterase activity and by calcium feedback to the rate of cGMP production. The model is governed by twelve parameters. By setting the model’s parameter values to match experimentally-derived values from cone or rod photoreceptors, the model can be configured to represent either photoreceptor type. For all primate retina modeling in this manuscript, we configured the photoreceptor model to represent primate rods by setting the initial values of the model parameters to those that were derived from separate patch-clamp experiments on primate rod photoreceptors (Chen et al. (2024); Supplementary Table. 1). For rat retina modeling, we configured the photoreceptor model as a cone photoreceptor for modeling responses at photopic light levels, and as a rod photoreceptor for modeling responses at scotopic light levels. The parameter values here were obtained from fitting the model to mouse cone and rod photoreceptors as part of patch-clamp experiments for other studies Chen et al. (2024). The corresponding values are stated in Supplementary Table. 1.

We implemented this biophysical model as a fully-trainable neural network layer, called the photoreceptor layer, using the Keras (Chollet et al., 2015) package in Python. All twelve parameters of the photoreceptor layer can be trained through backpropagation using the Keras and TensorFlow package in Python – although the user can also set some or all of these parameters to be non-trainable and hence held fixed at their initial value. Photoreceptor parameters were initialized to their known values (Supplementary Table 1). For the experiments presented herein, 7 of the parameters were set to be non-trainable. Some of these parameters like the concentration of cyclic guanosine monophosphate (cGMP) in darkness vary across rod and cone photoreceptor types (rods and cones). Other parameters governing cGMP conversion into current, calcium concentration in the dark, affinity for Ca^2+^, and hills coefficient are comparable across photoreceptor types. The remaining 5 parameters, set to be trainable or non-trainable depending on the model configuration, consisted of the photopigment decay rate *σ*, the phosphodiesterase (PDE) activation rate *η*, the PDE decay rate *ϕ*, the rate of Ca^2+^ extrusion *β* and *γ* that controls the overall sensitivity of the model to light inputs. These trainable parameters also differ across photoreceptor types. In the current version of the model, the photoreceptor parameters are shared by all the input pixels, and each pixel acts as an independent photoreceptor: I.e., the conversion of each pixel into photocurrents only depends on that pixel’s previous values and not on the values of the other pixels.

The photoreceptor layer converts each pixel of the input movie in units of receptor activations per photoreceptor per second (R*receptor^*−*1^s^*−*1^) into photocurrents (pA) by solving the differential equations using the Euler’s method. Similar to the conventional CNN model, the photoreceptor layer takes as input 80 frames, where each frame corresponds to 8 ms. The output of this layer is a movie that is 80 frames long, and the same spatial dimensions as the input visual stimuli. The first 20 frames of the photoreceptor layer output are truncated to account for edge effects. The photocurrents movie is then z-scored using Layer Norm. This normalization step is crucial due to substantial differences in scale between the biophysical model’s parameters and the downstream CNN weights. The absence of these normalization layers hinders the photoreceptor– CNN model’s convergence. The resulting movie is then passed through the downstream CNN layers, where the size of the first convolution layer filter representing the temporal dimension is 60 frames instead of the 80 frames in the case of conventional CNN model.

For modeling RGC responses across light levels (experiments of Figs. 4–7), the input to the photoreceptor layer was a movie segment of 180 frames. The first 60 frames were then discarded to account for edge effects. The longer movie segment allowed for longer integration times at the lowest light level of 0.3 R*receptor^*−*1^s^*−*1^.

### Linear photoreceptor–CNN architecture

The linear photoreceptor model consists of a linear convolutional filter given by Eq. (4), and described previously in Angueyra et al. (2022).

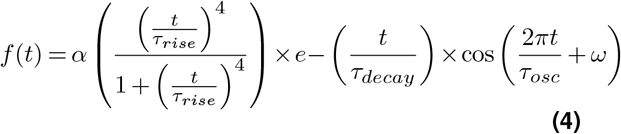

The parameters for this model were initialized to the following values: *α* = 631 pA/R*/s, *τ*_*rise*_ = 28.1 ms, *τ*_*decay*_ = 24.3 ms, *τ*_*osc*_ = 2 *×* 10^3^ s, and *ω* = 89.97 deg. These values corresponded to estimates of the single-photon response, obtained by recording cone photoreceptor responses (Angueyra et al., 2022). However, all the parameters were set to trainable and could therefore be learned along with the downstream CNN weights.

Similar to the other models, the hyperparameters of the linear photoreceptor–CNN model were optimized via a grid search.

### Model training

Model weights were optimized using Adam (Kingma and Ba, 2015), where the loss function was given by the negative log-likelihood under Poisson spike generation. The network layers were regularized with a *L*_2_ weight penalty at each layer, to prevent loss of information and be more robust to outliers. In addition, a *L*_1_ penalty was applied to the output of the fully-connected layer because the neural activity itself is relatively sparse and *L*_1_ penalties are known to induce sparsity. Learning rates were initially set to 0.001. A learning rate scheduler reduced the learning rate by a factor of 10 at epoch 3, 30 and 100.

The number of channels in each CNN layer and the filter sizes were optimized by a grid search for each model type and dataset. Grid search for each experiment and model type was conducted using the full training data for that experiment. During the grid search procedure, models were trained for 50 epochs. Optimal hyperparameters were selected by evaluating the model on validation data that was neither used during the training phase, or during the model evaluations of predicted responses. Models with these optimal hyperparameters were then re-trained for at least a 100 epochs.

### Model evaluation

Trained models were evaluated using the held out test dataset not seen during the training. We quantified the model performance with the fraction of explainable variance in each RGC’s response that was explained by the model (FEV). This quantity (Eq. (5)) was calculated as the ratio between the variance accounted for by the model and the *explainable* variance (denominator in Eq. (5)). Such metrics to quantify how well a model predicts neural data have been used in previous studies like Cadena et al. (2017). We calculate FEV as

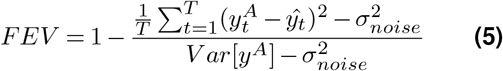

where,

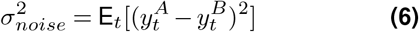

*y*^*A*^ and *y*^*B*^ are the observed spike rate of an RGC calculated as an average across set of repeats *A* and set of repeats *B* respectively. The sets *A* and *B* were obtained by randomly splitting the total number of repeats into two. ŷ_*t*_ represents the predicted spike rate by the model at time bin *t*. The explainable variance (denominator in Eq. (5)) is the variance of each RGC attributable to the stimulus, computed by subtracting an estimate of the observed noise from the variance across time (Eq. (7)) in the actual RGC’s responses, calculated as

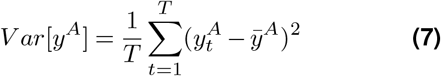

Where 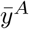 is the the observed spike rate *y*^*A*^ averaged across time. In all neural data sets we considered, the number of trials was sufficient (N = 10 for naturalistic movies, N = 225 for white noise movies) and hence the estimated noise variance was quite low. As a result, our FEV values are quite similar to what is obtained using the usual fraction explained variance calculation, which does not correct for unexplainable noise. By definition, FEV can be negative if the prediction error is larger than the variance in the actual responses. We report each model’s performance across all RGCs as the median FEV across the set of RGCs. For ease in interpretation, we present FEV as a percentage throughout our results.

### Model RGC temporal receptive fields (Fig. 6)

For a given model RGC, we computed the gradient of its output spiking rate with respect to the pixel values in the input movie segments, similar to Maheswaranathan et al. (2018); Goldin et al. (2022). These gradients were evaluated for different binary white noise movie segments from the primate retina experiment across light levels (Figs. 4,5). In total we had 400,000 input movie segments, generated by incrementing the white noise movie that was shown to the retina forward by one frame at a time (where 1 frame corresponds to 8 ms). These input movie segments spanned a total duration of 54 minutes. Since all the models were implemented with TensorFlow (Abadi et al., 2016), we calculated the gradients using automatic differentiation.

The resulting gradients matrix representing spatiotemporal receptive field were decomposed into spatial and temporal components (Supplementary Fig. 3) using Singular Value Decomposition (SVD), similar to the way the spatial and temporal receptive fields are computed from the spike triggered average (STA) analysis applied directly to experimental data.

We normalized the spatial component to have unit mean. By doing so, our process of decomposing the instantaneous spatio-temporal RF into spatial and temporal components assigned any variations in the receptive field’s amplitude only to the temporal component. The average of all the instantaneous temporal receptive fields was taken as the model RGC’s temporal receptive field. In Figs. 6c,d, the temporal receptive field was normalized by the maximum peak.

### Statistics and reproducibility

We used the SciPy package in Python to perform a two-tailed Wilcoxon signed-rank test to compare performance across models (Figs. 3, 5). This non-parametric test was chosen due to its appropriateness for paired samples (in this case the same RGCs being modeled by different architectures) and its robustness against potential violations of normality assumptions. The null hypothesis tested was that the difference between the median fraction of explainable variance explained (FEV) by the two models for the population of RGCs modeled was 0. In comparing response latencies between two different light levels obtained by different methods (Fig. 6e), we performed a two-tailed Wilcoxon rank sum test of the null hypothesis that there was no difference between the distributions (N = 22 RGCs) of latencies at the two different light levels.

## Data availability

All retina electrophysiology data used in this study can be made available upon reasonable request.

## Code availability

Codes for the proposed photoreceptor–CNN model and for the conventional CNN model used in this study are available at https://github.com/saadidrees/dynret.

## Acknowledgments

This work was supported by VISTA: Vision science to applications fellowship to SI; Canada Research Chair grant and NSERC Discovery grant (RGPIN-2019-06379) to JZ.

## Author contributions

All authors contributed to the design of the study. F.R. developed the biophysical photoreceptor model; S.I. implemented the biophysical photoreceptor model as a neural network layer and designed and implemented CNNs; J.Z. supervised the implementation of the biophysical photoreceptor model as a neural network layer and the design and implementation of CNNs; G.D.F. and M.B.M supervised retina electrophysiology experiments; J.Z. supervised the overall study. All authors wrote the manuscript.

