## Supplementary Information for "Biophysical neural adaptation mechanisms enable artificial neural networks to capture dynamic retinal computation"

[1]Department of Physics and Astronomy, York University, Toronto, ON M3J 1P3, Canada; [2]Centre for Vision Research, York University, Toronto, ON M3J 1P3, Canada; [3]Department of Ophthalmology, University of Washington, Seattle, WA 98195, U.S.A.; [4]Department of Physiology and Biophysics, University of Washington, Seattle, WA 98195, U.S.A.; [5]Stein Eye Institute, University of California, Los Angeles, CA 90095, U.S.A.; [6]Learning in Machines and Brains Program, Canadian Institute for Advanced Research, Toronto, ON M3J 1P3, Canada

### Supplementary Note 1. Biophysical model of phototransduction reproduced from Angueyra et al. (2022)

In the phototransduction cascade (Fig. ??c), continuous synthesis of cGMP by guanylate cyclase (GC) opens cGMP-gated channels in the membrane. Activation of light-sensitive opsin (Opsin*) results in channel closure through the activation of G-protein transducin (Gt*), subsequently activating PDE* and decreasing cGMP concentration. Calcium ions ($Ca^{2+}$) enter the photoreceptor outer segment via cGMP-gated channels and are extruded through $Na^{+}$ / $K^{+}$ / $Ca^{2+}$ exchangers in the membrane. These enzymatic reactions of the phototransduction cascade are represented as a set of six differential equations (reproduced below from Angueyra et al. (2022)).

Input to the model is the stimulus intensity as a function of time (in units of R*receptor$^{-1}$s$^{-1}$) given by $Stim(t)$ in Eq. S1. The output of the model is the photoreceptor outer segment current as a function of time, $I(t)$ given in Eq. S4.

In the first step of the model, the stimulus (Stim) activates opsin molecules (denoted as R for Receptor, R* when active), which decay with a rate constant $\sigma$ as follows:

$$\frac{dR^*(t)}{dt} = \gamma Stim(t) - \sigma R^*(t) \quad \textbf{(S1)}$$

Here, $\gamma$ is a scaling factor (or opsin gain factor) that controls the overall sensitivity of the model to light inputs.

Active opsin molecules then activate phosphodiesterase (PDE) molecules through transducin, so that the activity of PDE(P) is as follows:

$$\frac{dP(t)}{dt} = R^*(t) - \phi P(t) + \eta \quad \textbf{(S2)}$$

where $\phi$ is the decay rate constant of PDE, and $\eta$ is the PDE activity in darkness.

The concentration of cGMP in the outer segment (G) depends on the PDE-mediated hydrolysis and the rate of synthesis (S) by the guanylate cyclase (GC) as follows:

$$\frac{dG(t)}{dt} = S(t) - P(t)G(t) \quad \textbf{(S3)}$$

The outer segment current carried by the cGMP-gated channels depends on G and can be approximated as follows:

$$I(t) = k_{Ca} G(t)^h \quad \textbf{(S4)}$$

where $h$ denotes the effective cooperativity, and $k_{Ca}$ depends on the maximal current and the affinity of the channel for cGMP. $k_{Ca}$ is calcium dependent providing feedback to the cGMP-gated channels.

A fraction (q) of the outer segment current (I) is carried by calcium, so on exposure to light the calcium concentration (Ca) decreases. Calcium extrusion in the outer segment is mediated by the $Na^{+}$/$K^{+}$, $Ca^{2+}$ exchanger. This process is simplified in the model as a single expotential process with rate constant $\beta$ as follows:

$$\frac{dCa(t)}{dt} = qI(t) - \beta Ca(t) \quad \textbf{(S5)}$$

The calcium concentration regulates S (the rate of cGMP syntesis) following a Hill curve, as follows:

$$S(t) = \frac{S_{max}}{1 + \left(\frac{Ca(t)}{K_{GC}}\right)^m} \quad \textbf{(S6)}$$

where $S_{max}$ is the maximum synthesis rate, and $K_{GC}$ and m are the affinity and cooperativity constants.

A second feedback as a single-exponential process that is calcium dependent but as a smaller decay rate constant was modeled as follows:

$$\frac{dCa_{slow}(t)}{dt} = \beta_{slow}(Ca_{slow}(t) - Ca(t)) \quad \textbf{(S7)}$$

This value determines the value of $k_{Ca}$ as follows:

$$k_{Ca} = k \times \frac{1}{1 + \frac{Ca_{slow}}{Ca_{dark}}} \quad \textbf{(S8)}$$

Rapid adaptation emerges in this model emerges from changes in the rate of cGMP turnover produced by light-dependent changes in phospodiesterase activity and by calcium feedback to the rate of cGMP production.

In the proposed Keras photoreceptor layer, the parameters $\sigma$, $\gamma$, $\phi$, $\eta$, $q$, $\beta$, $C_{dark}$, $K_{GC}$, $\beta_{slow}$, $m$, $k_{Ca}$, and $h$ can all be set to trainable. At the start of model training, all parameters are set to their values determined through photoreceptor fits(Supplementary Table. 1). For primate retina models, we used parameter values reflecting primate rod photoreceptors (Chen et al., 2024). For rat retina model, we used mouse cone photoreceptor and rod photoreceptor values (Chen et al., 2024). The non-zero value of the slow calcium dependent feedback ($\beta_{slow}$) in the cone phototransduction model provided marginal gains when modeling rat responses at photopic light levels. We therefore set its value to 0 when training the model together with downstream CNN, effectively removing calcium feedback to cGMP-gated channels. This however did not affect the other calcium dependent feedback on the concentration of guanylate cyclase (GC) (Fig. **??**c).

Unless explicitly stated, we allowed the parameters $\sigma$, $\beta$, $\phi$ and $\eta$ and $\gamma$ to learn optimal values together with the downstream conventional CNN model through backpropogation.

| Parameter | Symbol | Units | Primate rod | Rat rod | Rat cone |
|---|---|---|---|---|---|
| Opsin gain | $\gamma$ | unitless | 4 | 9 | 10 |
| Opsin decay rate constant | $\sigma$ | s$^-1$ | 7.07 | 6.95 | 11.24 |
| PDE decay rate constant | $\phi$ | s$^-1$ | 7.07 | 6.95 | 11.24 |
| PDE dark activation rate | $\eta$ | s$^-1$ | 2.53 | 2.66 | 576 |
| cGMP-to-current constant | $k$ | pA$^2\mu$M$^{-3}$ | 0.01 | 0.01 | 0.01 |
| cGMP channel cooperativity | $h$ | unitless | 3 | 3 | 3 |
| $Ca^{2+}$ extrusion rate constant | $\beta$ | s$^-1$ | 25 | 25 | 3.47 |
| Channel feedback decay rate constant | $\beta_{slow}$ | s$^-1$ | 0 | 0 | 0.04 |
| $Ca^{2+}$ GC affinity | $K_{GC}$ | $\mu$M | 0.5 | 0.46 | 0.33 |
| $Ca^{2+}$ GC cooperativity | $m$ | unitless | 4 | 4 | 4 |
| $Ca^{2+}$ GC concentration in darkness | $Ca_{dark}$ | $\mu$M | 1 | 1 | 1 |
| cGMP concentration in darkness | cGMP$_{dark}$ | $\mu$M | 15.5 | 13.4 | 20 |

**Supplementary Table 1.** Parameters and best fit values for primate rod and mouse rod and cone phototransduction biophysical model.

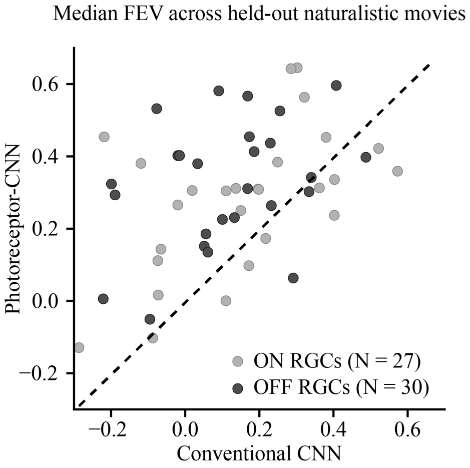


**Supplementary Figure 1. Photoreceptor–CNN model consistently outperforms conventional CNN model when cross-validated with different held-out naturalistic movies.** Conventional CNN and photoreceptor–CNN models originally trained on checkerboard movies were fine-tuned using data from different combinations of 8 naturalistic movies and evaluated using the held-out test movie. For different ON RGCs (light gray circles, N = 27) and OFF RGCs (dark gray circles, N = 30), we calculated the median FEV across the held-out naturalistic movies (N = 6 movies) for conventional CNN model (x-axis) and photoreceptor–CNN model (y-axis). Three movies which generated weak spiking activity in recorded RGCs were excluded from cross-validation.

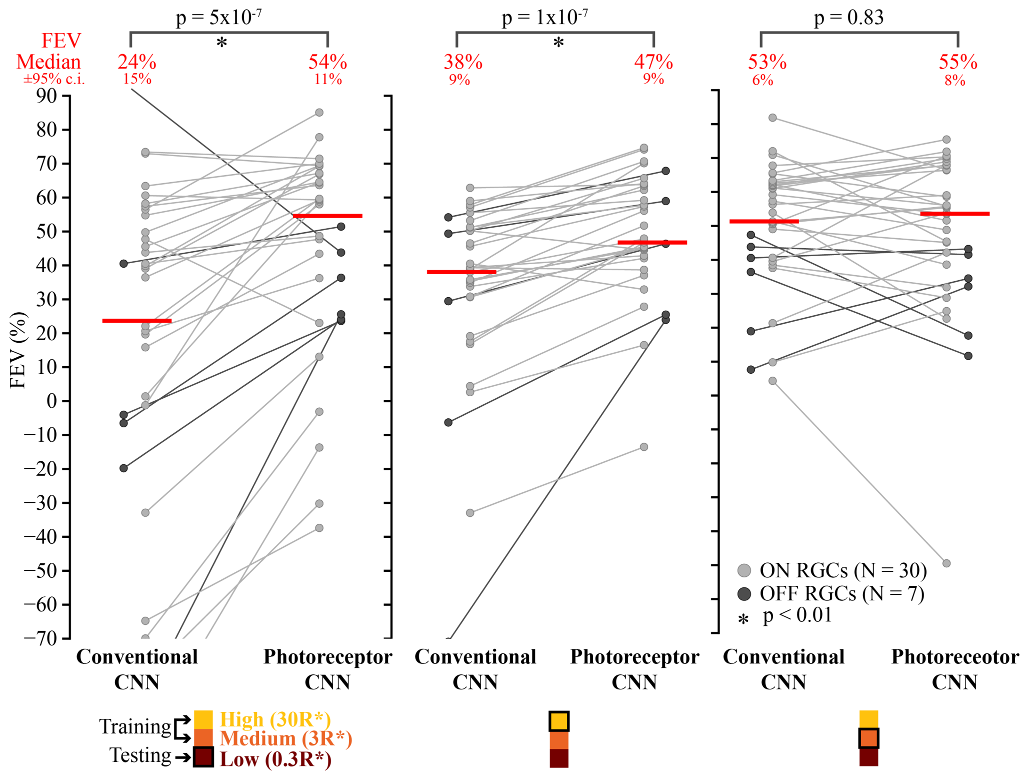


**Supplementary Figure 2.** Extended analysis showing photoreceptor-CNN model's superior performance at all combinations of training and test light levels of Fig. **??**c. For each panel, the legend below the box plot panel shows the two light levels the models were trained at and the third light level at which it was tested (black outline). The box plots show the distribution of FEVs at this testing light level. Testing light levels were low (left panel), high (middle panel), and medium (right panel). The y-axis shows the Fraction of Explainable Variance Explained (FEV) in each RGC's (circle) response by the conventional CNN (left) and the photoreceptor–CNN where the photoreceptor layer parameters were trained along with downstream CNN (right). Light gray circles denote ON type RGCs (N = 30), and dark gray circles denote OFF type RGCs (N = 7). Connecting lines in each panel link the FEV values for each RGC's performance across both models. Median FEV values across all RGCs (N = 37) are indicated by red lines, and stated as FEV $\pm$ $95\%$ $c.i.$ in red text. P-values were calculated by performing two-sample Wilcoxon signed-rank test on the FEV distributions from the CNN and photoreceptor-CNN model. An asterisk indicates statistical significance (p < 0.01) between performance of the two models.

### Supplementary Note 2. Photoreceptor–CNN model more accurately predicts context-dependent changes in RGC sensitivity

Frequent and significant changes in light intensity on the retina, such as those caused by saccades, alter RGC encoding (Idrees et al., 2020, 2022; Tikidji-Hamburyan et al., 2015; Ruda et al., 2020) and feature selectivity (Grimes et al., 2014). The use of CNNs to investigate such context-dependent changes in RGC feature selectivity is becoming increasingly useful (Maheswaranathan et al., 2018; Goldin et al., 2022; Walker et al., 2019). This is because instantaneous receptive fields of CNN-modeled RGCs can be calculated from gradients of the model-predicted response with respect to the input pixel values (Supplementary Figure 3). These derivatives can be evaluated for individual segments of input movie, without the need to average over multiple segments as would be needed in the typical reverse correlation analysis. As a result, the CNN-based methods provide a new tool for researchers to probe rapid stimulus-dependent changes in neuronal receptive fields.

We hypothesized that PR-CNN model could capture context-dependent changes in receptive fields better than conventional CNNs, due to the fact that it can better capture how the retina adapts to dynamically changing stimuli. Our focus was specifically on quantifying changes in RGC sensitivity, as defined by the gain (amplitude) of its receptive field.

To test the hypothesis, we computed the instantaneous spatial and temporal receptive fields (STRFs) of the model RGCs from gradients (Supplementary Fig. 4a) of the model trained on the primate retina dataset that was used to evaluate model performance across light levels (Figs. **??**,**??**). To obtain the instantaneous STRF of a given model RGC, we computed the gradient of its output spiking rate prediction with respect to the pixel values in the input movie segments, similar to Maheswaranathan et al. (2018); Goldin et al. (2022). These gradients were evaluated for different binary white noise movie segments from the experiment. In total we had 400,000 input movie segments, spanning a total duration of 54-minutes. Since all the models were implemented with TensorFlow, we calculated the gradients using automatic differentiation. We decomposed each resulting STRF into its spatial and temporal components using Singular Value Decomposition (SVD) and normalized the spatial component to have unit mean. This enabled us to represent the STRF's amplitude solely in the temporal component. We observed substantial variation in gain, i.e. the peak amplitude of these receptive fields varied across the movie segments for a given model RGC (Supplementary Figure 4a). This variation constitutes a model-derived prediction that the real RGCs (whose activities the model was trained to predict) would exhibit the same pattern of stimulus-dependent gain variation.

To test these model predictions, we sorted the movie segments by their model-predicted gains, and then divided the segments into 10 equally-sized bins (i.e., lowest-gain bin 1 up to highest-gain bin 10; Supplementary Figure 4b). We then estimated the RGC STRF separately based on the data in each bin, using the standard reverse correlation (Dayan and Abbott, 2001) approach. This procedure was performed separately for each RGC, because the stimulus segments that yield higher or lower gain can differ between RGCs. From these experimentally-derived STRFs, we then extracted the spatial and temporal components using SVD. The gains (peak of temporal filters) of these temporal receptive fields within each bin (Supplementary Figure. 4c) were consistent with model-derived predictions obtained from averaging the temporal receptive fields from the model's gradients (Supplementary Figure. 4b) over the set of stimuli within each bin. Thus, the retrospective analysis of RGC receptive fields in these stimulus bins confirmed that the photoreceptor–CNN model could veridically predict context-dependent RGC sensitivity.

To quantify the accuracy with which the model could predict RGC sensitivity changes, we calculated the squared error between the predicted (derived from the model gradients) and the actual (derived from reverse correlation) gain within each stimulus bin and averaged the resulting errors across bins. Supplementary Figure 4d shows a scatter plot of the resulting mean squared error (MSE) values for the photoreceptor–CNN and the conventional CNN model for different RGCs (N = 22) and different light levels (columns). These RGCs constitute a subset of the total RGCs that displayed FEV values greater than 50% at the high (30 R*receptor$^{-1}$s$^{-1}$) and medium (3 R*receptor$^{-1}$s$^{-1}$) training light levels for both models: thus, these were the RGCs for which both models' predictions were most reliable. The photoreceptor–CNN model consistently exhibited lower MSE values than the conventional CNN model at all light levels (Supplementary Figure. 4d-f) with the difference being statistically significant for medium (3 R*receptor$^{-1}$s$^{-1}$) light level (Supplementary Figure. 4e; p = 0.01, one-tail Wilcoxon signed-rank test, N = 22RGCs) and the held-out low test light level of 0.3 R*receptor$^{-1}$s$^{-1}$(Supplementary Figure. 4f; p = 0.007, one-tail Wilcoxon signed-rank test, N = 22 RGCs). This result indicates that the photoreceptor–CNN model is superior in predicting context-dependent changes in RGC sensitivity.

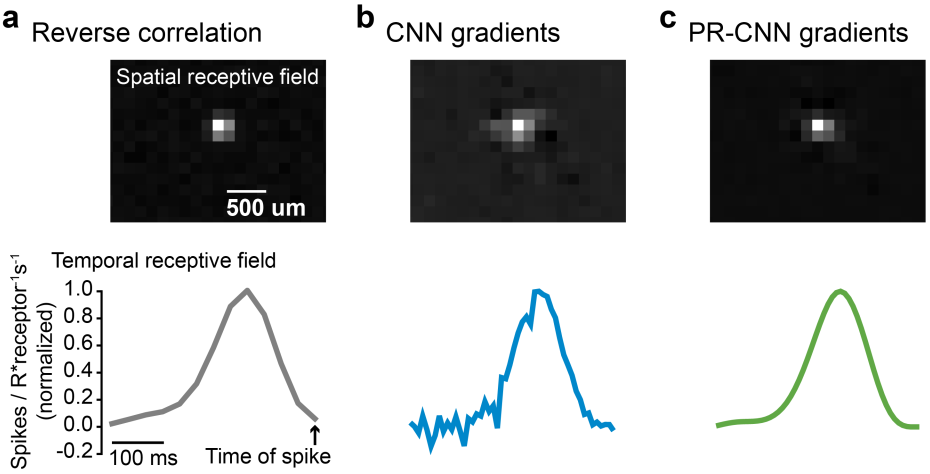


**Supplementary Figure 3. Receptive fields can be computed from model output gradients with respect to input pixel intensities. a.** The receptive field of an example RGC calculated from the reverse correlation of white noise movie ($\sim 55$ mins), decomposed into its spatial (top row) and temporal (bottom row) components. **b-c.** Spatial (top) and temporal (below) components of the receptive field of the same RGC calculated from the gradient outputs of the conventional CNN model, **b**, and the photoreceptor-CNN model, **c**, evaluated for a single movie segment ($\sim 1.5$ s).

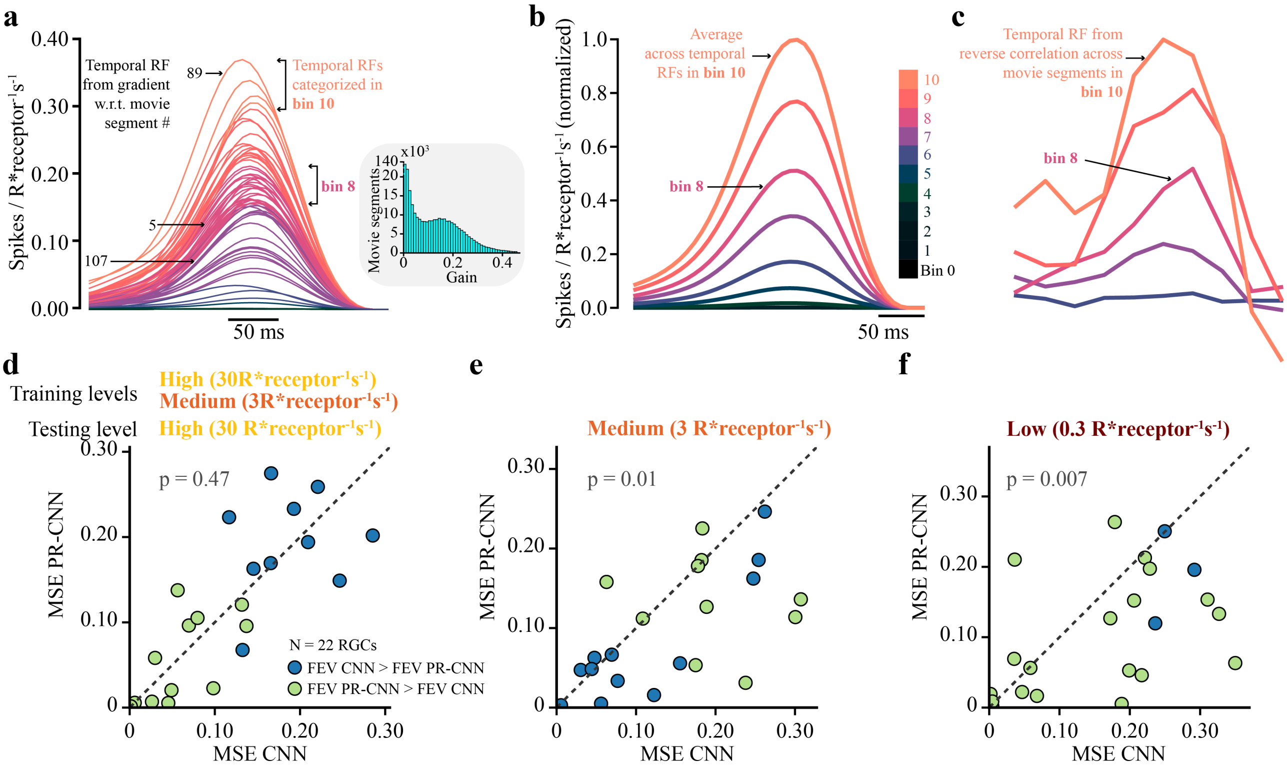


**Supplementary Figure 4. Temporal receptive fields estimated from photoreceptor–CNN model gradients better capture sensitivity changes across different input stimuli. a.** The temporal receptive fields of a model RGC obtained from gradients of the output of a PR-CNN model, evaluated for different movie segments (only a subset is shown here for clarity). Inset shows the distribution of receptive field gains obtained as the peak of the temporal filters across 400,000 individual movie segments. **b.** The temporal receptive fields in **a** were ranked by their gain, divided into 10 bins (different colors in **a**), and averaged within each bin. The averaged temporal receptive fields were normalized by the peak amplitude of the highest-amplitude curve. Color bar shows the line color for each bin. **c.** Temporal receptive fields calculated directly from reverse correlation on experimental data, using movie segments that were grouped into the same bins as in **b**. **d-f.** Mean squared error between the predicted gains, **b**, and actual gains, **c**, of the temporal receptive fields plotted for different RGCs (N=22) for conventional CNN model (x-axis) and PR-CNN model (y-axis). The models were trained at 30 R*receptor$^{-1}$s$^{-1}$and 3 R*receptor$^{-1}$s$^{-1}$, and predicted gains were obtained from gradients of model output w.r.t. movie segments at 30 R*receptor$^{-1}$s$^{-1}$**(d)**, 3 R*receptor$^{-1}$s$^{-1}$**(e)**, and 0.3 R*receptor$^{-1}$s$^{-1}$**(f)**. p-values were obtained by conducting a one-tailed Wilcoxon signed-rank test to analyze whether the PR-CNN MSE values for the population of RGCs were lower than those of CNN.

### Supplementary Note 3. Photoreceptor–CNN model training procedure for generalization across photopic and scotopic light levels (Fig. ??)

The photoreceptor–CNN experiments modeling rat RGC responses (Fig. **??**) was performed as follows. First, we trained the model end-to-end (including photoreceptor layer parameters) to predict rat RGC responses from Retina A (N = 61 RGCs) to stimuli at the bright (photopic) light level (Step 1 in Fig. **??**f). This led to an estimate of photoreceptor and CNN parameters for the cone pathway. We then estimated parameters reflecting rod photoreceptors by re-training the model at the dim (scotopic) light level, while keeping the CNN weights and biases fixed to their already-learnt values (Step 2 in Fig. **??**f). Thus, the retraining altered only the photoreceptor parameters based on the data from Retina A.

We then trained the photoreceptor–CNN model to predict RGC responses of another rat retina (Retina B) to stimuli at the bright light level (Fig. **??**f). Here, we fixed the photoreceptor parameters to the values already learned from Retina A at the bright light level. We then exchanged those photoreceptor parameters with the ones learned from Retina A at the low light level, while keeping the remaining CNN parameters fixed (Step 4 in Fig. **??**f). In this manner, we took the model where the CNN weights were learned on model of Retina B at the bright light level, and simply exchanged the cone photoreceptor parameters for rod parameters learned from retina A. Notably, in this procedure, none of the data from Retina B at the dim light level was used in training the model: these data were used only in evaluating the generalization ability of the photoreceptor–CNN model.
